# LsrL modulates Lsr2-induced chromatin structure to tune biosynthetic gene cluster regulation in *Streptomyces venezuelae*

**DOI:** 10.1101/2025.11.23.690019

**Authors:** Vivian Ramirez, Lauren Tiller, Marcin Jan Szafran, Xiafei Zhang, Dagmara Jakimowicz, Marie Elliot, Lydia Freddolino

## Abstract

Specialized biosynthetic gene clusters in *Streptomyces* are subject to complex regulation involving both transcriptional control and chromosome organization. The nucleoid-associated protein Lsr2 silences many of these clusters, yet how it shapes the global chromatin structure and how its conserved paralog LsrL contributes to this process remain poorly understood. In this study, we applied a multi-omics approach, combining transcriptional activity, genome-wide protein-DNA binding profiles, and three-dimensional chromosome conformation to characterize the coordination of Lsr2 and LsrL in exerting transcriptional control and genome architecture in *Streptomyces venezuelae*. In line with established Lsr2 functions, we find that Lsr2 sets broad transcriptional boundaries, while LsrL acts in a more context-specific manner that depends on the presence of Lsr2 and may function to reinforce or modulate Lsr2-mediated silencing. Loss of Lsr2 reshaped the chromatin landscape genome-wide, relieving its restriction on short-range contacts, triggering strong transcriptional changes and new domain boundaries near de-repressed biosynthetic gene clusters. These findings establish Lsr2 as a dominant but contextually modulated regulator whose interplay with LsrL coordinates specialized metabolism with higher-order chromosome organization.

## Introduction

Across bacterial genomes, nucleoid-associated proteins are highly abundant proteins that shape chromosome structure while altering gene expression [1–6]. These functions are especially vital throughout the complex life cycle of the soil-dwelling *Streptomyces* bacteria [3,7,8]. In *Streptomyces*, nucleoid-associated proteins not only drive nucleoid compaction but also coordinate gene regulatory networks that are essential for progression through their distinct life cycle stages [7,9–12]. *Streptomyces* are also renowned for their abundance of biosynthetic gene clusters (BGCs), each of which encodes the proteins needed to synthesize cluster-specific specialized metabolites. These metabolites are bioactive natural products that are presumed to provide a competitive advantage against other organisms that *Streptomyces* encounter in their native environments [13,14]. Since the discovery of these metabolites in the 1940s, many of these natural products have been harnessed for medicinal and agricultural application, including many widely used antibiotics [15,16]. Any given *Streptomyces* genome can contain anywhere between 8 and 83 BGCs [17–19]. A recently published study identified a total of 440 known BGC families (i.e., those showing high similarity to experimentally characterized clusters in the MIBiG database) and 8,799 unknown families from an analysis of 2,371 *Streptomyces* genomes, demonstrating that much of the metabolic potential of this genus remains to be explored [20]. However, the products of these BGCs are often cryptic and their expression remains repressed under standard laboratory conditions, making it difficult to investigate their metabolic potential [21,22].

Unlike most bacteria, *Streptomyces* possess a linear chromosome that is organized into distinct regions including a core region (which houses essential genes important for cell viability), and left and right terminal domains (also referred to as chromosomal arms, which house a large number of genes that are poorly conserved across *Streptomyces,* and notably include many BGCs) [23–26]. During metabolic differentiation when specialized metabolism initiates, the linear chromosome undergoes conformational changes where the chromosome adopts a closed conformation (with the two terminal arms coming into close physical proximity) and the terminal regions become transcriptionally active [27]. While these conformational transitions were reported in *Streptomyces ambofaciens*, which did not undergo sporulation under the conditions tested, *Streptomyces venezuelae* has been shown to undergo large-scale chromosomal rearrangement during sporulation [7]. Such dynamic transitions highlight the necessity of regulatory mechanisms to ensure precise control over the activation and repression of chromatin in the terminal regions.

Among the nucleoid-associated proteins responsible for the repression of horizontally-acquired genes is Lsr2, a protein that is conserved across the Actinomycetes, and is functionally analogous to H-NS in the enterobacteria [28–30]. Lsr2 preferentially binds to AT-rich sequences within the genome, where it can oligomerize to form long nucleoprotein filaments and/or bridge distant segments of the DNA, which both facilitates compaction of the chromosome and represses transcription by obstructing RNA polymerase (RNApol) activity [29–31]. Although much of the early research into Lsr2 focused on its role in the mycobacteria, where it functions as a xenogenic silencer and regulates many virulence-related genes [29], recent work has expanded to include its role in organizing and regulating the *Streptomyces* genome. Within *S. venezuelae*, Lsr2 binds to a majority of predicted BGCs throughout the genome, repressing their expression and effectively serving as a metabolic gatekeeper [32,33]. A prime example of this is the chloramphenicol BGC, where Lsr2 mediates transcriptional repression by binding to sites within and upstream of the cluster, and both polymerizes along the DNA and bridges these disparate segments, ultimately limiting the production of chloramphenicol [31]. While many global and pathway-specific regulators of BGCs are well characterized [31,34–36], the extent to which Lsr2 cooperates with other nucleoid-associated proteins in controlling chromosomal structure and gene expression remains unclear.

In addition to Lsr2, *Streptomyces* species also encode an Lsr2-like paralog, known as LsrL. However, little is known about whether LsrL functions independently or in association with Lsr2. In this study, we used an integrative genomic approach, combining RNA-seq, RNApol ChIP-seq, global protein occupancy profiling, and Hi-C to define the shared and unique transcriptional targets of Lsr2 and LsrL and to examine their influence on chromosome organization. We show that Lsr2 acts as the primary silencer of genes within and beyond BGCs, exerting broad effects on transcription and chromosomal structure. In contrast, LsrL regulates a smaller, more select set of genes and can function either antagonistically or in a supplementary role depending on the genomic context. Our findings demonstrate an asymmetrical regulatory relationship between Lsr2 and LsrL, where LsrL functions in a predominantly Lsr2-dependent manner and where loss of LsrL results in continued, though partial, repression stemming from Lsr2 compensation. Genome-wide chromatin state analysis revealed Lsr2-regulated states that shape nucleoid organization and transcriptional activity in the terminal chromosomal regions. By integrating Hi-C data, we reveal a role for Lsr2 in constraining short-range chromosomal contacts throughout the genome and suppressing domain boundary formation near highly repressed BGCs. Together, these findings highlight the complex interplay between Lsr2 and LsrL in coordinating specialized metabolism with higher-order chromosome architecture in *S. venezuelae*.

## Results

### LsrL contributes to region-specific modulation, while Lsr2 enforces predominant repression at key loci

Lsr2’s role in repressing specialized metabolic clusters, particularly the chloramphenicol BGC, is well established in *S. venezuelae* [32]. However, the contribution of LsrL to this regulatory landscape is largely uncharacterized. To begin dissecting their respective influences, we first examined representative genomic regions, including known Lsr2 targets, to assess how deleting *lsr2*, *lsrL*, or both impacted transcription and genome-wide protein occupancy. To this end, we took advantage of previously constructed single and double knockout mutants of *lsr2* and/or *lsrL* in *S. venezuelae* and conduRNActed RNA-seq on these mutants, alongside wildtype (WT) cells grown in MYM liquid medium for 16 hours, corresponding to the pre-sporulation phase, before large-scale chromosomal compaction occurs [7]. We complemented these transcriptional data with *in vivo* protein occupancy display at high resolution (IPOD-HR) and RNApol chromatin immunoprecipitation sequencing (RNApol ChIP-seq), which together profile both global protein-DNA interactions and RNApol binding. In the IPOD-HR approach, total protein occupancy is measured genome-wide, and the RNApol signal is subtracted (using the RNApol ChIP-seq data collected in parallel) to quantify occupancy by other regulatory and architectural proteins [37]. Both IPOD-HR and RNApol ChIP-seq samples were briefly treated with rifampicin prior to fixation, which inhibits promoter clearance but not elongation, thereby enriching RNApol signal at active promoters. To contextualize our data with known Lsr2 binding sites, we also reprocessed previously published Lsr2 ChIP-seq datasets [32] and visualized them alongside our occupancy and transcriptomic profiles.

We re-scanned the *S. venezuelae* chromosome for BGCs using antiSMASH [38] and identified 32 BGCs (two more than predicted by Gehrke *et al*. (2019) [32]; **Supplementary Table 1**). Among them, the chloramphenicol BGC spans over 41 genes (as predicted by AntiSMASH [38]) and serves as a well-established target of Lsr2-mediated silencing [31,32]. While antiSMASH may overestimate BGC boundaries, we used its annotations for consistency across all predicted clusters. At a representative subset of genes in the chloramphenicol BGC (*vnz_04455*–*vnz_04490*), we observed a reduction in protein occupancy in the Δ*lsr2* and Δ*lsr2*Δ*lsrL* mutants (as observed by IPOD-HR), alongside increased RNApol occupancy and increased transcription when compared to WT and the Δ*lsrL* mutant (**Figure 1A, D**). The presence of strong Lsr2 ChIP-seq signals at this site (observed using previous data [32]) further confirmed direct Lsr2 binding, supporting the interpretation that the observed loss of protein occupancy reflects, at least in part, the removal of Lsr2. However, given that IPOD-HR captures total protein occupancy, we cannot rule out contributions from other DNA-binding proteins in the WT strain, including LsrL. Indeed, deleting *lsrL* also triggered changes in transcription at the chloramphenicol BGC. Somewhat paradoxically, RNApol occupancy was slightly reduced in the Δ*lsrL* strain compared to WT, but several genes within the cluster also showed increased transcript abundance, with log2 fold changes exceeding 1 but remaining well below those observed in the Δ*lsr2* or Δ*lsr2*Δ*lsrL* mutants; it is not clear whether these effects are a result of transcriptional or post-transcriptional (possibly indirect) effects of LsrL at this locus. We also found that relatively high protein occupancy was maintained in the Δ*lsrL* mutant upstream of *vnz_04475* (*cmlS*), suggesting that Lsr2 could compensate for the absence of LsrL by continuing to bind and repress this locus. This pattern suggested that Lsr2 may continue to partially repress transcription in the absence of LsrL, consistent with the persistent protein occupancy signal observed in Δ*lsrL*. Taken together, these results indicate that while Lsr2 remains the dominant repressor of the chloramphenicol BGC, LsrL provides a supplementary regulatory role, with loss of LsrL causing partial de-repression of chloramphenicol biosynthetic genes.

**Figure 1.**
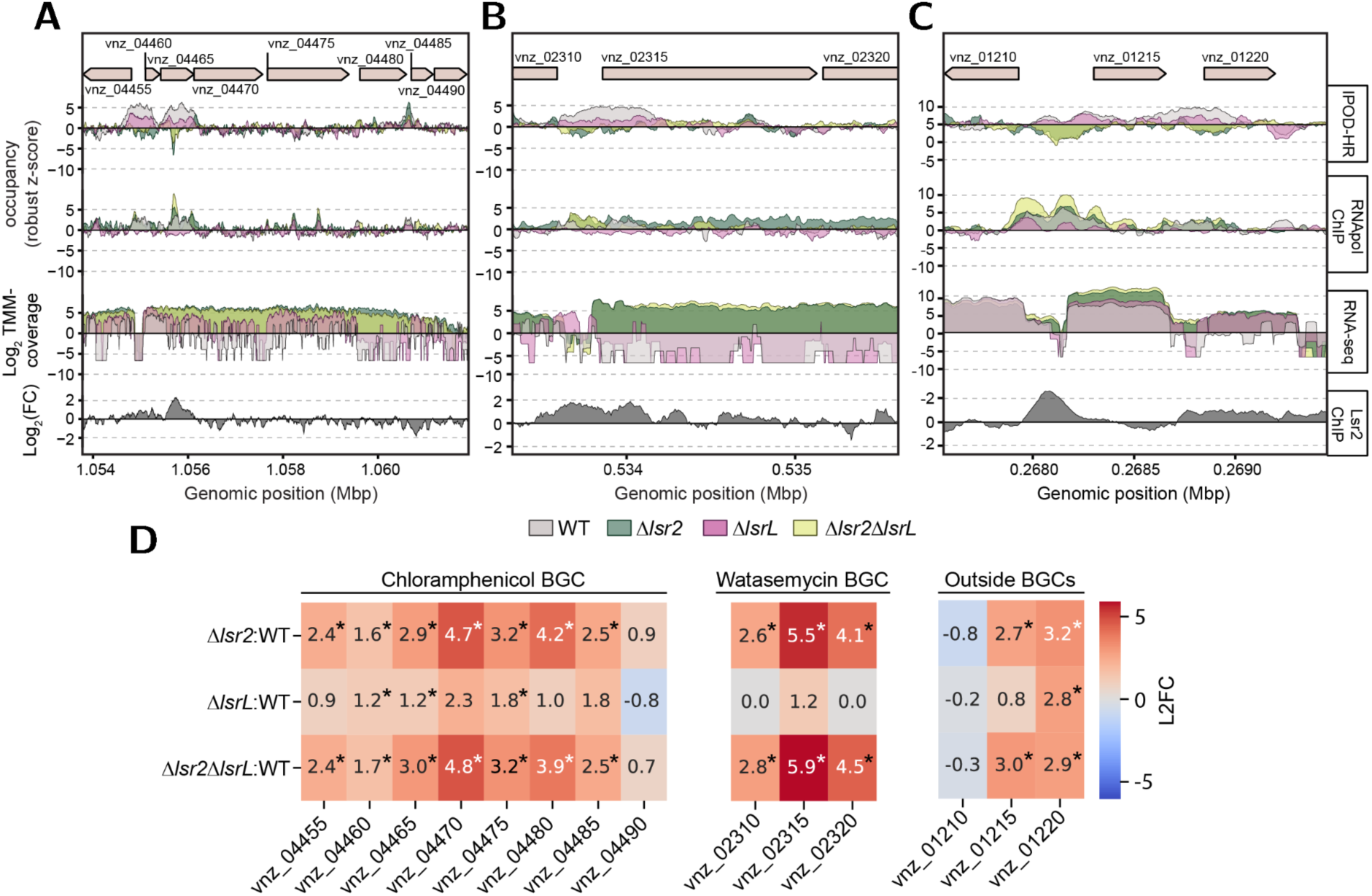
Chromatin and transcriptional profiles within and outside of BGCs reveal a co-regulatory relationship between Lsr2 and LsrL. Changes in RNApol and IPOD-HR occupancy (represented as robust z-scores relative to input), RNA-seq read coverage (represented as Log2 TMM-normalized coverage relative to genome-wide trimmed mean), and Lsr2 occupancy (represented as Log2 fold change associated with 3×FLAG tagged Lsr2 relative to a non-FLAG tagged Lsr2 variant) are shown. Profiles depicted are for (**A**) the chloramphenicol BGC, (**B**) the watasemycin BGC, and (**C**) a non-BGC locus, across WT (gray), Δ*lsr2* (green), Δ*lsrL* (pink), and Δ*lsr2*Δ*lsrL* (yellow) genotypes. BGCs from panels A-B were identified based on antiSMASH [38] predictions. Lsr2 ChIP-seq data were obtained from Gehrke et al. (2019) [32]. (**D**) Log2 fold changes (L2FCs) in gene expression for genes within the chloramphenicol BGC, watasemycin BGC, and genes outside BGCs in the Δ*lsr2*, Δ*lsrL*, and Δ*lsr2*Δ*lsrL* mutants relative to WT. Genes significantly up- or down-regulated are indicated by asterisk (|L2FC| ≥ 1 and q-value < 0.05).

To determine whether this regulatory profile was unique to the chloramphenicol BGC, or extended to other BGCs and potentially beyond – we explored additional examples of Lsr2-mediated regulation across BGCs, based on antiSMASH predictions, where there was a clear Lsr2 binding site (in the WT). One such example, encodes the biosynthetic machinery responsible for synthesizing watasemycin or a closely related specialized metabolite (*vnz_02310*–*vnz*_*02320*). Here, we observed a similar profile to that of the chloramphenicol BGC: in the Δ*lsr2* and Δ*lsr2*Δ*lsrL* mutants, we observed decreased global protein occupancy, coupled with increased RNApol occupancy and transcript abundance (**Figure 1B, D**). Unlike for the chloramphenicol cluster; however, loss of LsrL did not significantly impact either transcript levels or RNApol occupancy, with only one of the three genes exhibiting a modest increase in gene expression (**Figure 1D**), suggesting that this cluster was primarily controlled by Lsr2.

We observed a different pattern of genetic control by Lsr2 and LsrL at three genes shown in **Figure 1C** (not associated with a BGC), encompassing genes encoding a cupin domain-containing protein (*vnz_01210*), a SPW repeat protein (*vnz_01215*), and a DUF805 domain-containing protein (*vnz_01220*) and where Lsr2 binding was confirmed. Here, *vnz_01220* appeared to be cooperatively repressed by both Lsr2 and LsrL, as indicated by an equally-large increase in RNA read coverage in the Δ*lsr2*, Δ*lsrL*, and Δ*lsr2*Δ*lsrL* mutants. This AND-logic, where both Lsr2 and LsrL may be required to maintain repression and loss of either results in derepression, is in contrast to the independent repression by Lsr2 alone observed for the watasemycin BGC, highlighting the diverse mechanisms by which Lsr2 and LsrL regulate transcription. Overall, these findings underscore the dominant silencing role of Lsr2 while indicating a more nuanced regulatory function for LsrL.

### The terminal regions of the S. venezuelae chromosome exhibit distinct protein occupancy and transcriptional activity relative to the core

The left and right arms of the *Streptomyces* chromosome harbor a high density of BGCs and horizontally acquired genes, which can be subject to complex regulation and transcriptional silencing [7,23,25,26,31,32,39]. To explore whether the localized patterns of occupancy observed extend to broader chromosomal regions, we calculated 20 kb rolling averages of global protein occupancy and RNApol occupancy across the genome in the WT strain collected at 16 hours (pre-sporulation stage). Over the length of the chromosome, the IPOD-HR and RNApol ChIP-seq data revealed distinct protein occupancy patterns in the core versus the arm regions (right terminal domain (RTD) and left terminal domain (LTD)) of the chromosome (**Figure 2A**). The core region (spanning 1.9-6.3 Mbp) [7] was characterized by higher average RNApol occupancy and lower average occupancy by DNA-binding proteins, compared with the left and right terminal domains, which exhibited a reciprocal profile, with lower average RNApol occupancy and higher average global protein occupancy. Notably, in the absence of Lsr2 and Lsr2/LsrL the average RNApol occupancy increased in the terminal regions relative to WT, further supporting a role for Lsr2 in repressing transcriptional activity in these regions (**Supplementary Figure 1A**). Interestingly, a subregion within the LTD of the WT strain (∼1-1.9 Mbp) displayed occupancy patterns more similar to the core region than to the rest of the LTD (**Figure 2A**), suggesting that regulatory or structural distinctions may exist within the LTD itself. Overall, these data suggest greater transcriptional activity in the core region relative to the terminal regions under the physiological condition examined here. In contrast, the elevated protein occupancy in the terminal regions suggests a higher level of binding by nucleoid-associated proteins including Lsr2 and LsrL, likely shaping the chromatin architecture. However, a comprehensive understanding of how these protein-DNA interaction landscapes contribute to broader chromatin states and their regulatory implications in the context of Lsr2 and LsrL, requires a more integrative framework that combines occupancy and expression data over time and across genotypes.

**Figure 2.**
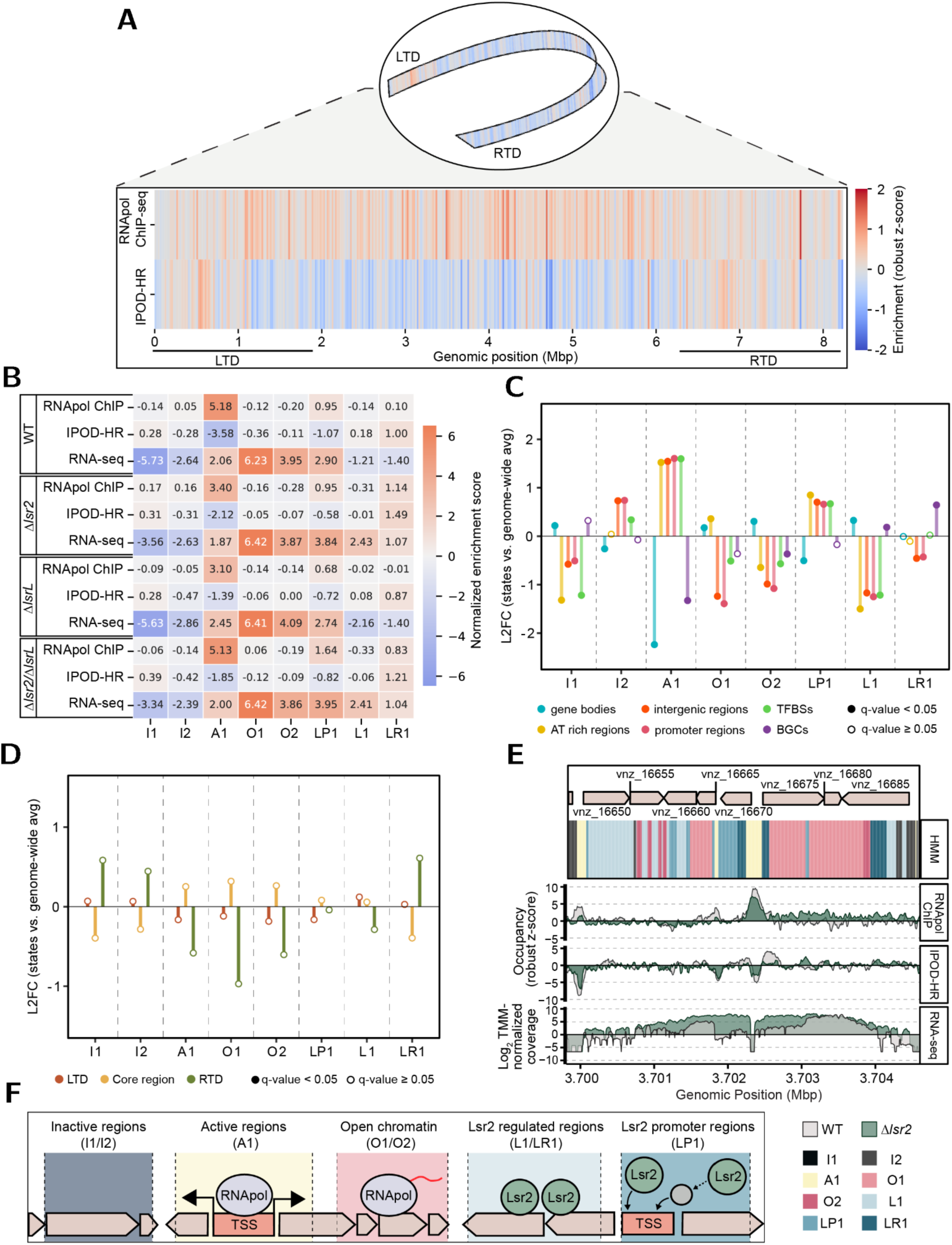
Protein occupancy in WT strain highlights regulatory patterns, with genotype-dependent chromatin state changes revealing diverse regulatory landscapes that align with genomic features. (**A**) Heatmap of protein occupancy derived from RNApol ChIP-seq and IPOD-HR data in WT cells. Enrichment is represented as robust z-scores (relative to an input sample) that were calculated as rolling averages in 20 kb windows. The left terminal domain (LTD) and right terminal domain (RTD), separated by the core region (∼1.9-6.3 Mbp), are indicated. (**B**) Averages of the normalized enrichment scores across RNApol ChIP-seq, IPOD-HR, and RNA-seq data for WT, Δ*lsr2*, Δ*lsrL*, and Δ*lsr2*Δ*lsrL* within assigned chromatin states containing regions characterized as inactive (I1, I2), active (A1), open chromatin (O1-O2), Lsr2-regulated promoters (LP1), and Lsr2-regulated regions (L1, LR1). (**C**) L2FC of the proportion of regions within each chromatin state relative to the genome-wide average for genomic features (gene bodies, TFBSs, BGCs, AT-rich, intergenic, and promoter regions). (**D**) L2FC of chromatin state proportions relative to the genome-wide average, as in panel C, but grouped by chromosomal regions (LTD, core, and RTD). In panels C-D, lollipop heads indicate statistical significance from an approximate permutation test (10,000 iterations; filled: q-value < 0.05, unfilled: q-value ≥ 0.05). (**E**) Chromatin state changes in a representative snapshot of the core region of the chromosome illustrating changes in RNApol and IPOD-HR occupancy (represented as robust z-scores relative to input), and RNA abundance (represented as Log2 TMM-normalized coverage relative to genome-wide trimmed mean) in Δ*lsr2* (green) and WT (gray) strains. (**F**) Schematic representation of chromatin state classifications inferred from HMM analysis. Inactive regions (I1, I2) exhibit little to no protein occupancy. Active regions (A1) are characterized by RNApol occupancy at promoter regions, including transcription-start sites (TSSs) and active transcription. Open chromatin regions (O1, O2) exhibit pervasive transcription across gene bodies. Lsr2 regulated regions (L1, LR1) span genes with increased transcription in the absence of Lsr2. Lsr2 promoter region (LP1) includes promoter regions with increased transcription in the absence of Lsr2.

### Chromatin state modeling reveals the genome-wide chromatin composition of transcriptional regulation and Lsr2-associated repression

To provide a framework for understanding the regulatory roles of Lsr2 and LsrL genome-wide, we characterized chromatin states across the genome using a Hidden Markov Model (HMM), an approach commonly used in studies of eukaryotic epigenomic regulation and chromatin structure [40,41]. We integrated the IPOD-HR, RNApol ChIP-seq, and RNA-seq data from the WT, Δ*lsr2*, Δ*lsrL*, and Δ*lsr2*Δ*lsrL* mutants to train a Gaussian HMM (GHMM) using normalized enrichment scores, enabling the inference of hidden (chromatin) states associated with the observed occupancy and gene expression patterns across the genome at a 5 bp resolution.

To determine the optimal number of states for the GHMM, we performed a 5-fold cross-validation using the genomic data partitioned across the genome. The GHMM was trained using a window size of 1,678 bp, chosen to capture the varying sizes of RNApol ChIP-seq and IPOD-HR enrichment peaks while allowing for convenient division of the chromosome. As noted above, the protein occupancy profiles for IPOD-HR and RNApol ChIP-seq were obtained after brief treatment with rifampicin, which restricts RNApol signal to active promoters; accordingly the RNA-seq data reflects the resulting transcriptional output, whereas RNApol ChIP-seq represents promoter-proximal occupancy. Using cross-validated mean absolute error to evaluate the model performance, we found that eight chromatin states were optimal for modeling the genomic data (**Supplementary Figure 1B**). To better understand the transcriptional landscapes defined by the chromatin state model, we manually categorized the eight states into five broad qualitative categories based on their average enrichment scores across genotypes and their associated genomic features: inactive regions (I1–I2), active promoter regions (A1), open chromatin (O1–O2), Lsr2-regulated regions (L1–LR1), and Lsr2-regulated promoters (LP1; **Figure 2B-C**). While these five categories reflect overarching functional trends, each of the eight states remains distinct and was analyzed independently throughout.

The I1 and I2 chromatin states reflected transcriptionally inactive regions, with low average enrichment scores for RNApol ChIP-seq, IPOD-HR and RNA-seq abundance.

However, I2 regions were, on average, less repressed than I1, as observed by their slightly higher (less negative) RNA-seq abundance. These differences were echoed in their genomic feature composition, where I1 was enriched for gene bodies and depleted of transcription factor binding sites (TFBSs) and AT-rich regions relative to the genome-wide average, whereas I2 regions showed enrichment for intergenic regions, promoter regions, and TFBSs. This suggested that I2 may encompass inactive regulatory regions, while I1 broadly reflects silent gene bodies.

The A1 chromatin state exhibited enhanced average enrichment of RNApol and RNA-seq read abundance, and negative IPOD-HR occupancy across all genotypes, which is consistent with actively transcribed areas. These regions likely encompassed transcriptional initiation sites, given the RNApol enrichment observed relative to the other states, combined with the effects of rifampicin noted above, where RNApol was engaged in transcription with few regulatory proteins bound. This was further supported by the enrichment of this state in intergenic regions, promoter regions, TFBSs, and AT-rich regions, while being depleted in gene bodies and BGCs.

The O1 and O2 chromatin states, categorized as “open chromatin”, were characterized by an enrichment of RNA-seq read abundance and a depletion in RNApol and IPOD-HR average enrichment scores, across all genotypes. Both O1 and O2 states were depleted in intergenic regions, promoter regions, and TFBSs, but were enriched in gene bodies. The “open” chromatin regions thus appear to correspond to actively transcribed regions outside of the promoters themselves (*e.g.*, gene bodies of currently expressed genes). Notably, O2 exhibited higher levels of AT-richness relative to O1, which may reflect differences in genomic context or suggest that these open regions are associated with functionally distinct sets of genes.

The LP1 chromatin state regions showed higher transcriptional activity in the absence of Lsr2, as reflected by both the RNA-seq and RNApol ChIP-seq data, when compared to the WT and Δ*lsrL* mutant. This pattern suggested that while these regions may already be somewhat transcriptionally active, Lsr2 further limits their expression, and its absence results in transcriptional derepression. The elevated average RNApol occupancy specifically in the Lsr2-deficient mutants indicates that Lsr2 may restrict RNApol binding or progression at the regulatory regions of these loci. Furthermore, the observation that these regions were enriched for promoter regions, intergenic regions, TFBSs, and AT-rich regions while being depleted in gene bodies, further reinforced this hypothesis.

The L1 chromatin state regions had low occupancy by RNApol and regulatory proteins across genotypes, but showed pronounced increases in transcript levels stemming from the loss of Lsr2, and decreased transcript levels as a result of loss of LsrL. L1 was enriched for gene bodies and BGCs and was depleted for intergenic regions, promoter regions, TFBSs, and AT-rich regions. This likely reflected a chromatin structure associated with Lsr2-repressed genes (many of which may additionally reflect cases where LsrL antagonized Lsr2-mediation repression).

LR1 chromatin state regions were characterized by moderate protein occupancy across all genotypes, with a slight increase in protein occupancy in the Δ*lsr2* and Δ*lsr2*Δ*lsrL* mutants relative to WT and Δ*lsrL*, suggesting compensatory binding by other regulatory proteins at these loci upon loss of Lsr2. These regions also showed elevated RNApol and RNA-seq enrichment specifically in the Δ*lsr2* and Δ*lsr2*Δ*lsrL* mutants, indicating that Lsr2 acts as a transcriptional repressor at these loci. Unlike L1, LR1 regions appeared to undergo a more robust transcriptional activation, with both increased polymerase activity and altered regulatory protein binding. In addition, these regions were depleted in intergenic and promoter regions but showed enrichment for BGCs, aligning with the observation that these states capture Lsr2-mediated repression within BGC-encoded loci.

Given that Lsr2 silences horizontally acquired genes that are enriched in the terminal domains of *S. venezuelae* [32], and that Lsr2 deletion led to increased RNApol occupancy in these regions (**Supplementary Figure 1A**), we examined chromatin state variation across the chromosome (**Figure 2D**) and tested for differences in the overall rates of occurrence of each chromatin state in the major regions. Although no chromatin state reached statistical significance, the RTD showed slight enrichment for I1, I2, and LR1 states and depletion of A1, O1, O2, and L1 states. In contrast, the LTD showed chromatin state frequencies largely consistent with genome-wide distributions. The lack of statistical significance associated with the RTD chromatin states was likely due to the large size of chromosomal regions relative to individual chromatin states (5 bp resolution), with small shifts in state proportions being diluted when analyzed across broad regions. Nevertheless, these trends were consistent with a role for Lsr2 as a xenogenic silencer, primarily repressing transcription in the right terminal arm.

Meanwhile, we observed that the core region remained accessible to transcriptional machinery, supporting active gene expression (**Figure 2A, D**).

To examine how chromatin states transition within a given window we focused our attention on the eight-gene region spanning *vnz_16650*–*vnz_16685* within the chromosomal core (**Figure 2E**), as this locus exemplified the local chromatin state transitions captured by the GHMM, providing a visually clear instance of changes among the inferred states. The products of these genes included a MarR transcriptional regulator (*vnz_16655*), a PIN domain-containing protein (*vnz_16660*), a helix-turn-helix transcriptional regulator (*vnz_16675*), and a DUF397 domain-containing protein (*vnz_16680*). Consistent with our broader findings (**Figure 2A-C**), promoter regions with increased RNApol occupancy were classified as A1 states, while regions with elevated protein occupancy and/or increased RNA read abundance in the absence of *lsr2* (relative to WT) were designated as L1, LP1 or LR1. Areas exhibiting high RNA read levels in both WT and Δ*lsr2*, but lacking increased RNApol occupancy were assigned as O1 or O2 states, with highest abundance levels assigned to O2. The region encompassing genes *vnz_16650* through *vnz_16670* showed strong changes in transcription and protein occupancy upon *lsr2* deletion, and were predominantly assigned to the Lsr2-dependent chromatin states (L1, LP1, and LR1). In contrast, regions upstream of *vnz_16650* and *vnz_16685* (relative to each gene’s transcription start site) appeared to be Lsr2-independent, as indicated by the absence of RNA read abundance and RNApol occupancy in Δ*lsr2* and were therefore classified as I2 chromatin state. Overall, the observed changes in chromatin states, along with differences in genomic features between states, particularly in the Lsr2-regulated states relative to others, point to the potential for Lsr2 to modulate local and global chromatin landscapes in orchestrating gene regulation; we have schematized our description of the Lsr2-associated regulatory states in **Figure 2F**.

### LsrL has a more targeted regulon, and functions in the context of Lsr2

Having defined chromatin states based on protein occupancy and transcription, we next examined how these states corresponded to genome-wide expression patterns, with the goal of characterizing the global regulatory role of LsrL in relation to that of Lsr2. Loss of *lsr2* had a dramatic effect on the transcriptome, with 1,392 genes significantly upregulated and 892 genes significantly downregulated relative to WT (|log2 Fold Change (L2FC)| ≥ 1, q-value < 0.05; **Figure 3A**). Of these, 15.6% were associated with specialized metabolism (as predicted by antiSMASH [38]). In contrast, the *lsrL* mutant showed a much more modest effect, with 363 genes being significantly upregulated and 383 significantly downregulated, 14.9% of which were encoded in BGCs (**Figure 3B**). A highly similar expression profile to that of Δ*lsr2* was observed for the double Δ*lsr2*Δ*lsrL* mutant, with 1,277 genes being significantly upregulated and 886 being significantly downregulated when compared to WT (**Figure 3C**). When examining the significantly altered gene expression changes across chromosomal regions, both Δ*lsr2* and Δ*lsr2*Δ*lsrL* exhibited the greatest number of extremely upregulated genes in the LTD and RTD of the genome (L2FC ≥ 5, q-value < 0.05; **Supplementary Figure 2A, C**). In these regions, the distribution of L2FC values was strongly right-skewed, reflecting a disproportionate number of genes that were significantly upregulated, whereas the core region displayed a more symmetric distribution. In contrast, most genes in the Δ*lsrL* mutant were downregulated in the RTD and upregulated in the core/LTD, hinting at a possible dual role for LsrL in activating a larger number of genes in the RTD and repressing a smaller set of genes in the core and LTD (**Supplementary Figure 2B**). Overall, these findings confirm a broad repressive role for Lsr2, as had been reported in Gehrke *et al*. (2019) [32]. In contrast, LsrL appears to contribute to the control of fewer genes. Notably, there was a significant increase in *lsrL* expression in Δ*lsr2* (L2FC = +0.58, q-value = 0.03), while *lsr2* expression remained unaffected in the absence of LsrL. Despite the increased levels of *lsrL* transcripts in Δ*lsr2*, it is noteworthy that the transcriptional landscape of the Δ*lsr2*Δ*lsrL* most closely resembled that of Δ*lsr2*, suggesting that in Δ*lsr2*, either increased *lsrL* transcription did not lead to increased protein levels, or there were minimal effects of increased LsrL protein levels in a Δ*lsr2* background.

**Figure 3.**
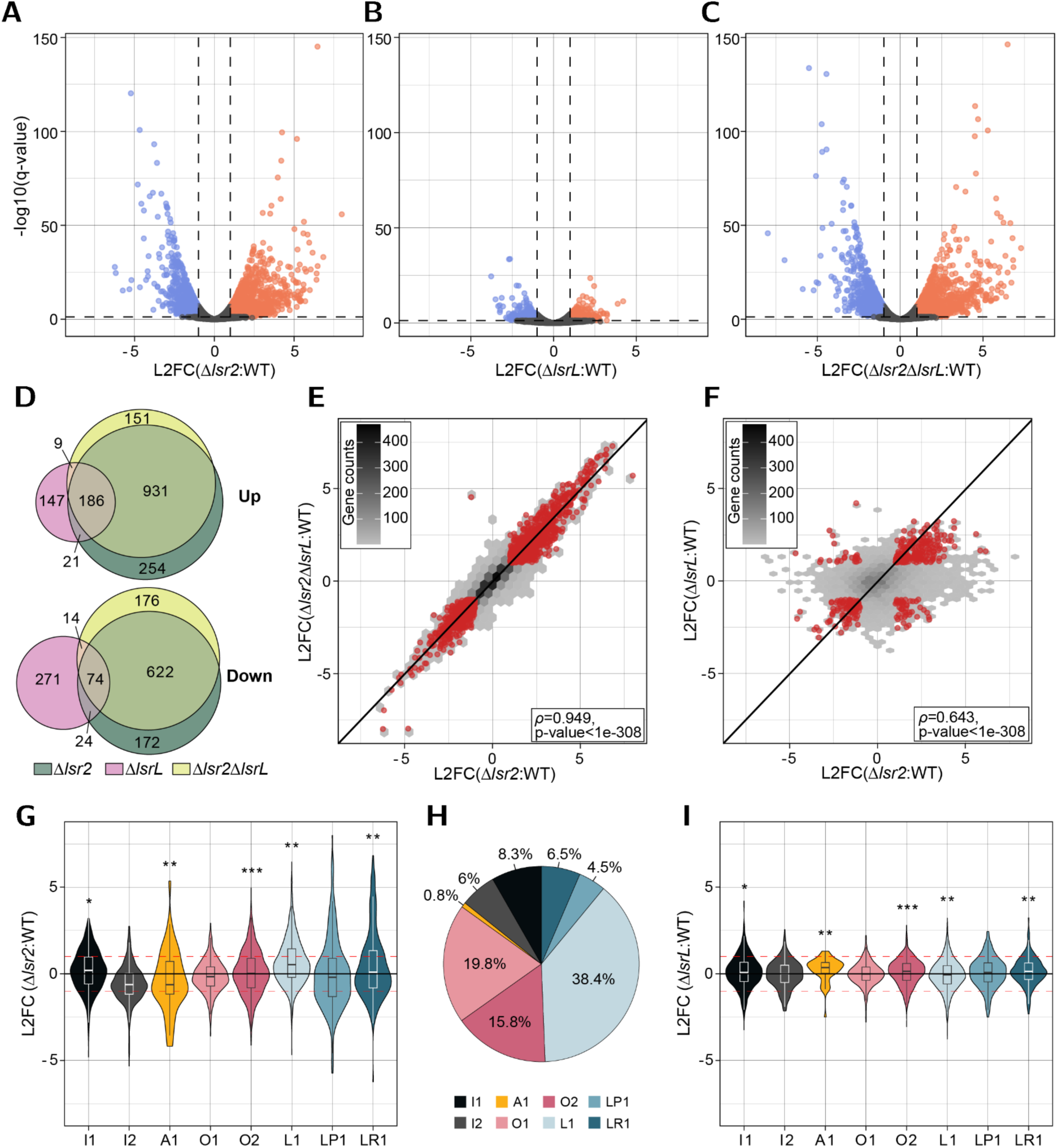
Lsr2 functions as a global transcriptional repressor whereas LsrL targets a more defined gene set. (**A-C**) Volcano plots of differentially expressed genes in the (**A**) Δ*lsr2*, (**B**) Δ*lsrL*, and (**C**) Δ*lsr2*Δ*lsrL* mutants relative to WT. Significantly upregulated (orange) and significantly downregulated (blue) genes are defined as those with |L2FC| ≥ 1 and q-value < 0.05. (**D**) The comparison of unique and shared genes that were significantly up- or downregulated in the Δ*lsr2* (green), Δ*lsrL* mutant (pink), and Δ*lsr2*Δ*lsrL* (yellow) mutants across panels A-C. The number of significantly altered genes in the (**E**) Δ*lsr2* vs. Δ*lsr2*Δ*lsrL* mutants and (**F**) Δ*lsr2* vs.Δ*lsrL* mutants (relative to WT) are shown. The hexbin plots represent the gene density, ranging from black (highest gene counts) to gray (lowest gene counts) bins. Genes overlapping between mutants that are significantly up- or downregulated are highlighted in red. Spearman correlations coefficients were calculated and are displayed on each plot. (**G**, **I**) L2FCs in gene expression for genes grouped by chromatin states in the (**G**) Δ*lsr2* and (**I**) Δ*lsrL* mutants relative to WT. Boxplots exhibit the median and interquartile range within each distribution. Statistical significance (q-value) was computed using a Wilcoxon signed-rank test with Benjamini-Hochberg FDR correction and is denoted by asterisks. Red dashed lines indicate L2FC ± 1. (**H**) The proportion of genes assigned to each chromatin state corresponding to the genes whose L2FC distributions are shown in panels G and I.

To identify specific genes differentially regulated by Lsr2 and LsrL (either directly or indirectly), we compared the significantly altered genes between the genotypes. Comparing gene expression profiles in Δ*lsr2* and Δ*lsr2*Δ*lsrL* (each relative to WT) revealed a strong correlation in differentially expressed genes (𝜌 = 0.949; p-value < 1e-308), with 1,813 genes showing consistent expression profiles across both genotypes (L2FC| ≥ 1, q-value < 0.05; **Figure 3D, E**). Among them, 1,117 (61.6%) were significantly upregulated and 696 (38.4%) were significantly downregulated in the mutants. The Δ*lsr2* and Δ*lsrL* single mutants showed a weaker correlation (𝜌 = 0.643; p-value < 1e-308), with only 305 genes similarly regulated, most of which were significantly upregulated (207 upregulated, 98 downregulated; **Figure 3D, F**). The notable similarity of the Δ*lsr2*Δ*lsrL* expression profile to that of Δ*lsr2*, suggests that rather than an additive effect of the individual *lsr2* and *lsrL* deletions, the regulatory effects of LsrL may depend on the presence of Lsr2. However, the opposing expression changes observed for a subset of genes between Δ*lsr2* and Δ*lsrL* (**Figure 3F**) further indicate that LsrL can act antagonistically to Lsr2 at certain loci, promoting transcription where Lsr2 exerts repression and vice versa.

Given the distinct differential expression profiles of the Δ*lsr2* and Δ*lsrL* single mutants, we performed an information-theoretic pathway analysis (using the iPAGE [42] software) for gene set enrichment to further explore the biological pathways regulated by each factor (**Supplementary Figure 2D-E**). In the absence of *lsr2*, we observed systematic increases in expression for several GO terms associated with biosynthesis and metabolism (*e.g.*, *de novo* inosine monophosphate (IMP), arginine, and chorismate biosynthesis), energy production (such as ATP synthesis coupled electron transport and cytochrome c-oxidase activity), molecular binding and transport activities (including phospholipase C, cation transport, and NADH dehydrogenase), as well as viral-related processes characteristic of prophages (**Supplementary Figure 2D**). Conversely, terms showing systematic decreases in expression in the absence of *lsr2* were primarily associated with core elements of gene expression, including nucleic acid-protein interactions such as DNA and RNA binding, along with core gene expression processes (*e.g.*, transcription initiation, translation). Deleting *lsrL* led to the enrichment of similar GO terms as for Δ*lsr2*, including those related to energy production, molecular binding and transport activity, and biosynthetic and metabolic processes (**Supplementary Figure 2E**). However, unlike in Δ*lsr2*, genes associated with transcription and translation-related GO terms, including translation and ribosomal subunits, were generally increased in expression (rather than decreased) in the absence of *lsrL*. The enrichment of biosynthetic, metabolic, and energy production processes in both Δ*lsr2* and Δ*lsrL* mutants suggests that each protein participates in regulating core metabolic and energy pathways.

However, Lsr2 may have a broader regulatory role in controlling endogenous foreign DNA elements, as indicated by the enrichment of viral-related processes in genes increasing in transcription when *lsr2* is deleted.

Building on the chromatin state classifications defined in **Figure 2**, we explored whether the regulatory impact of Lsr2 and LsrL varied across these states. Lsr2 had the largest effect on gene expression within regions classified as L1 and LR1, which were states observed to have strong Lsr2-mediated regulation (**Figure 3G**). Consistent with our assignment, genes in the L1 state, which encompassed over 38% of all coding sequences, showed an average L2FC of 1.54 in the Δ*lsr2* relative to WT (**Figure 3H**). In contrast, LsrL had a minimal regulatory influence across all chromatin states (**Figure 3I)**. While statistical tests detected significant deviations from zero in Δ*lsrL*, the median shifts were close to zero and were well within the interquartile range, suggesting a negligible biological impact. Taken together, these results reinforce a predominant role for Lsr2 in shaping the global transcriptome and chromosome organization, including across chromatin states that differ in genomic features.

### BGC-encoded gene expression profiles reveal coordinated regulation by Lsr2 and LsrL

Prompted by previous findings by Gehrke *et al*. (2019) [32] that the transcription of specialized metabolic genes in BGCs (and associated metabolite production) were significantly impacted by Lsr2, we next investigated the overlap between different types of BGCs and the changes in gene regulation revealed by our datasets. Using the 32 BGCs predicted by antiSMASH [38], we identified 24 unique BGC types based on their classification. We cross-referenced each BGC type with the differential expression information from our RNA-seq experiments, applying a L2FC threshold of ± 1 and a q-value < 0.05 (**Figure 4**). Across the genes encoded within BGCs, loss of *lsr2* resulted in 222 significantly upregulated and 112 significantly downregulated genes, relative to WT (**Figure 4A**). Genes affected by *lsr2* deletion primarily included those within BGCs that encoded polyketide synthases (PKS) and non-ribosomal peptide synthetases (NRPSs; **Figure 4A**). Notably, genes encoding machinery responsible for synthesizing ectoine represented the only BGC class that lacked any genes significantly upregulated in Δ*lsr2*. In contrast, Δ*lsrL* had a more stringent response, with only 47 genes significantly upregulated and 64 significantly downregulated amongst those encoded in BGCs, when compared to WT (**Figure 4B**). The types of BGCs showing the largest changes in the absence of *lsrL* included ectoine, lanthipeptide class II/terpene, NRPS, and PKS. Of the genes encoded in BGCs, 18.6% of those significantly impacted by Lsr2 and 55.9% of those significantly affected by LsrL were co-regulated by both proteins (n=62), with most showing similar directionality in their associated expression changes, apart from the ectoine biosynthetic cluster (**Figure 4C**). Consistent with our genome-wide gene expression profile observations, the differentially expressed BGC-encoded genes were largely similar in the Δ*lsr2* and Δ*lsr2*Δ*lsrL* mutants, with 279 genes showing similar expression changes in both strains; these genes represented over 99% of the total BGC-encoded genes affected in these mutants. NRPS, PKS, and lanthipeptide-encoding clusters were among the major BGC classes that had the greatest number of shared genes between Δ*lsr2* and Δ*lsr2*Δ*lsrL* mutants. Indeed, there was only one gene (*vnz_02580*) from the lanthipeptide class, encoding an MFS transporter, that showed a differential regulatory effect in the single versus double mutant (**Figure 4D**), where read counts mapping to *vnz_02580* were low in the Δ*lsr2* mutant (similar to WT) but were substantially higher in the Δ*lsr2*Δ*lsrL* mutant. Overall, these results confirm a pre-eminent role for Lsr2 in regulating genes within BGCs, where it largely suppresses gene expression. In contrast, deleting *lsrL* had a more subtle effect, with the majority of those genes impacted by the loss of LsrL being shared with Lsr2 (55.9%).

**Figure 4.**
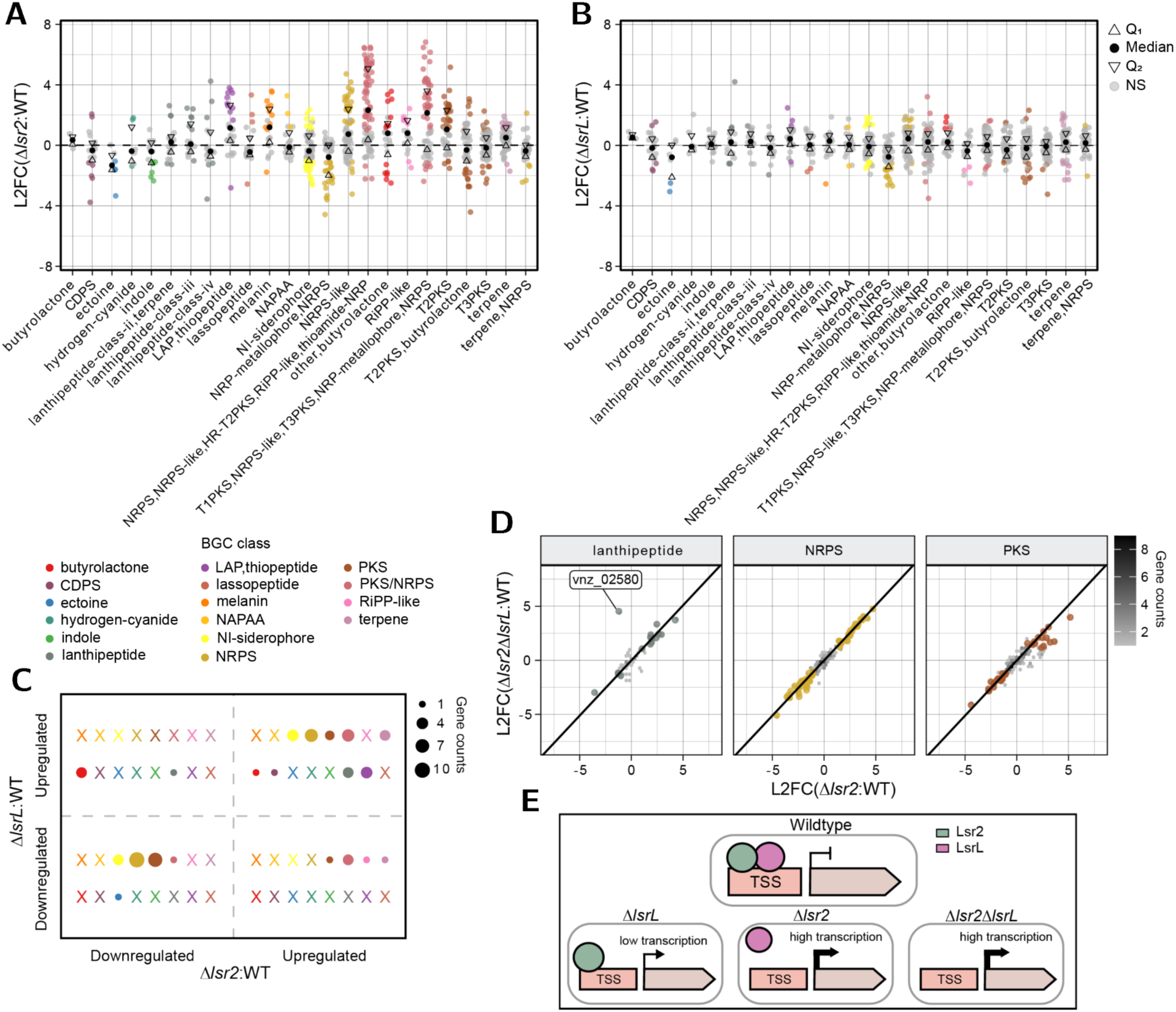
Lsr2 has broad impacts on multiple BGC classes, whereas LsrL effects are more constrained and are largely Lsr2-dependent. (**A-B**) L2FCs in gene expression across BGC classes (predicted by antiSMASH) in Δ*lsr2* (**A**) and Δ*lsrL* (**B**) mutants relative to WT. Significantly upregulated and downregulated genes are highlighted in their corresponding BGC class (|L2FC| ≥ 1, q-value < 0.05). First quartile (Q_1_, upward triangles), median (black circles), and third quartile (Q_3_, downward triangles) values are displayed. Genes not significantly (NS) regulated are displayed in gray. (**C**) Quadrant map of the significantly overlapping upregulated or downregulated gene targets (n=60) within BGC classes in Δ*lsrL* versus Δ*lsr2* mutants relative to WT (|L2FC| ≥ 1, q-value < 0.05). Quadrants display an X for BGC classes having no overlapping significant genes between mutants. Data point size denotes the gene count within each BGC class. Gene distribution: upper right quadrant (n=29), upper left quadrant (n=4), lower left quadrant (n=23), and lower right quadrant (n=6). (**D**) Comparison of L2FCs in the Δ*lsr2* and Δ*lsr2*Δ*lsrL* mutants within the lanthipeptide, NRPS, and PKS BGC classes relative to WT. Genes significantly up- or downregulated in both mutants (relative to WT) are colored accordingly (|L2FC| ≥ 1, q-value < 0.05), with hexbin plots showing gene density. The single labeled gene in the lanthipeptide class (RS02595=MFS transporter) was oppositely regulated, showing increased expression in the Δ*lsr2*Δ*lsrL* mutant, while decreased expression in Δ*lsr2* (Δ*lsr2*: L2FC=-2.26, q-value=8.86e-07; Δ*lsr2*Δ*lsrL*: L2FC=2.76; q-value=1.43e-09). BGC class abbreviations: RiPP-like, other unspecified ribosomally synthesized and post-translationally modified peptide product; PKS, polyketide synthase; NRPS, non-ribosomal peptide synthetase; NI-siderophore, NRPS-independent siderophore; NAPAA, non-alpha poly-amino acids; LAP-thiopeptide, linear azol(in)e-containing peptides-thiopeptide; CDPS, tRNA-dependent cyclodipeptide synthases. (**E**) Proposed model of Lsr2–LsrL coordinated regulation at loci where both proteins target the same site. In WT, Lsr2 and LsrL bind transcription start sites (TSSs) to repress transcription. In the Δ*lsrL* mutant, Lsr2 remains bound and maintains repression, but less effectively, resulting in a modest increase in transcription relative to WT. In the Δ*lsr2* mutant, LsrL is unable to bind, leading to derepression and a large transcriptional increase relative to WT. In the Δ*lsr2*Δ*lsrL* mutant, the absence of both proteins results in derepression, with transcript levels similar to the Δ*lsr2* single mutant.

Together, with our global expression analysis and chromatin state modeling, the similarity in gene expression profiles between the Δ*lsr2* single mutant and Δ*lsr2*Δ*lsrL* double mutant, coupled with the lack of an additive effect when *lsrL* was lost from the Δ*lsr2* mutant, suggested that LsrL acts largely in an Lsr2-dependent manner (**Figure 4E**). Specifically, in the absence of Lsr2, LsrL appears unable to exert its regulatory function. In contrast, the persistence in protein occupancy and moderate expression changes in the Δ*lsrL* mutant indicate that Lsr2 may partially compensate for the loss of LsrL, although less effectively. Lastly the observation that LsrL co-regulates a fraction of Lsr2 regulated genes, coupled with the fact that Lsr2 represses *lsrL* expression, supports a role for LsrL in fine-tuning gene expression within specific genomic loci.

### Higher-order chromosome architecture appears to be largely shaped by Lsr2

The capacity of Lsr2 to bind and bridge DNA within and between adjacent regions [31] led us to examine whether its effects, along with those of LsrL, contributed to chromosome organization at higher levels. To address this, we employed Hi-C to map DNA contact frequency across the *S. venezuelae* chromosome (**Figure 5A, Supplementary Figure 3A-B**). In the WT strain, Hi-C contact maps exhibited a strong primary diagonal corresponding to interactions between neighboring loci and a secondary diagonal resulting from interarm contacts- both of which were consistent with earlier findings [7]. Using a similar analytical approach [7], principal component 1 derived from a principal component analysis of normalized Hi-C contacts, recapitulated the terminal regions, with LTD spanning ∼0-1.9 Mbp and RTD spanning ∼5.9-8.2 Mbp in the WT strain (**Supplementary Figure 4A**). Loss of Lsr2 in the Δ*lsr2* and Δ*lsr2*Δ*lsrL* mutants resulted in increased short-range contacts between neighboring loci in the core and terminal regions (**Figure 5A**). This trend is reflected in the contact decay curves where *lsr2* mutants maintain higher normalized contacts at short genomic distances than WT and Δ*lsrL* (**Supplementary Figure 3C**). Notably, a pronounced accumulation of DNA contacts in cells lacking *lsr2* was particularly evident near ∼4 Mbp, suggesting a large rearrangement of the local chromosomal organization in the core region, near the origin of replication. The Δ*lsr2* and Δ*lsr2*Δ*lsrL* mutants also exhibited a decrease in long-range interdomain contacts when compared to the WT strain. In contrast, deleting *lsrL* alone resulted in a moderate disruption in both short-range and long-range contacts, indicating a more limited structural perturbation across the chromosome (**Figure 5A**). Although the overall distributions of absolute differential contacts were qualitatively similar between Δ*lsr2* and Δ*lsrL*, the moderate Spearman correlation (⍴ = 0.55, p-value < 1e-324) indicated that these changes occurred at partially overlapping but distinct genomic regions (**Supplemental Figure 4B, C**). Taken together, Lsr2 exerts a broad impact on higher-order chromosomal architecture, while the LsrL effects are more nuanced.

**Figure 5.**
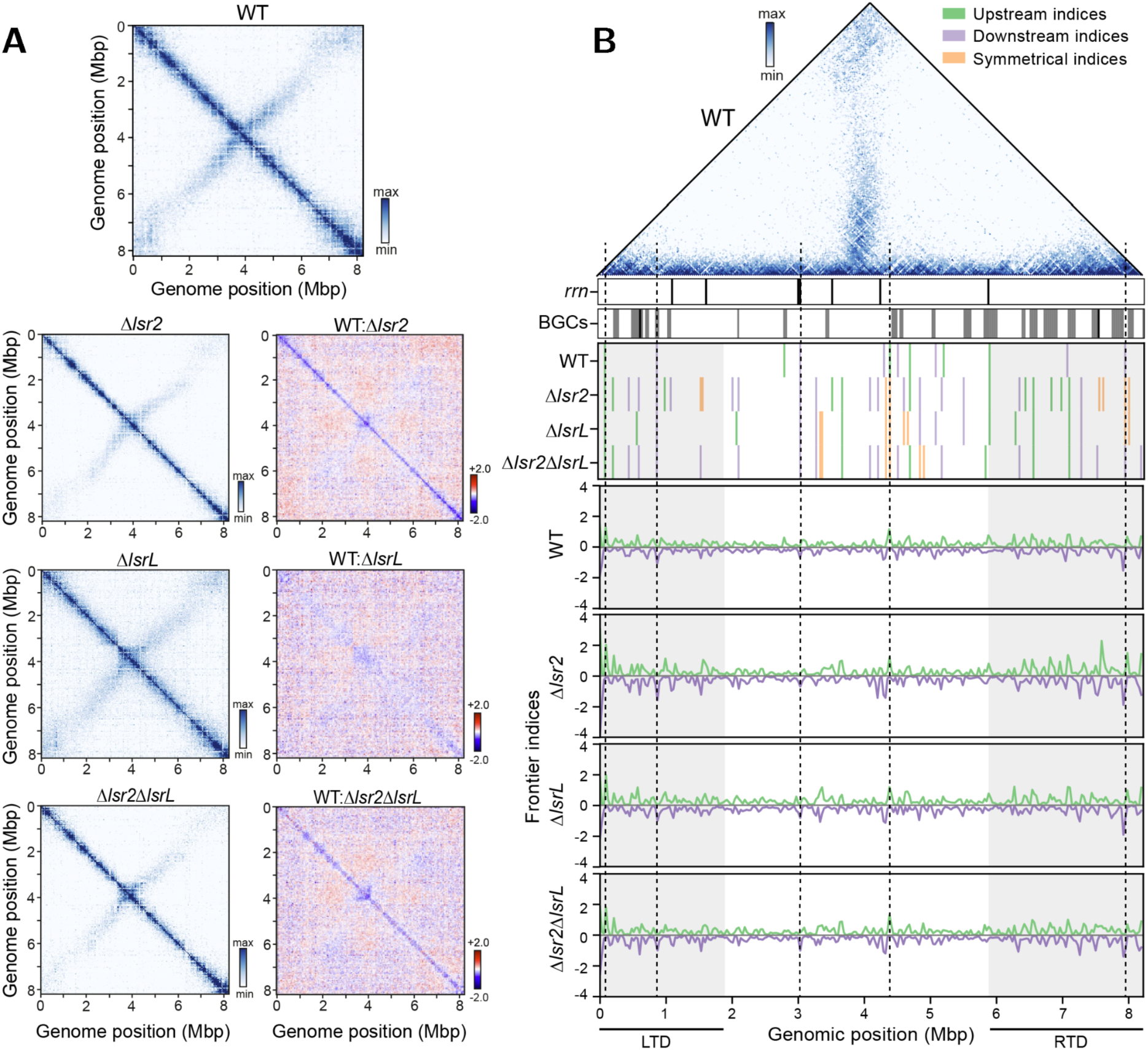
Deletion of Lsr2 and LsrL increases local chromosome interactions and promotes boundary formation. (**A**) Left: The normalized Hi-C contact map obtained for the WT (top), Δ*lsr2*, Δ*lsrL*, and Δ*lsr2*Δ*lsrL* mutant strains (replicate 1). Right: differential Hi-C maps in the logarithmic scale (log2) comparing the contact enrichment in WT relative to Δ*lsr2*, Δ*lsrL*, or Δ*lsr2*Δ*lsrL* mutant strains. The chromosomal coordinates were binned at a 30 kb resolution along both axes. (**B**) CID boundary detection using frontier index analysis [27,64], which identifies the change in contact bias between a genomic bin and its neighboring regions. A CID boundary is defined as a bin with a significant change in upstream contact bias, downstream contact bias, or a symmetrical shift in both directions (upstream and downstream peaks within ± 2 bins), relative to a WT threshold. Row 1: normalized Hi-C contact map for the WT strain shown as an upper triangle heatmap. Row 2: annotation of ribosomal operon locations across the chromosome. Row 3: annotation of BGCs encoded across the chromosome. Row 4: heatmap of significant frontier index peaks across genotypes. Peaks are classified as upstream-only (green), downstream-only (purple) or symmetrical (orange), and are considered significant if they exceed the WT median plus two standard deviations, calculated separately for LTD (0-1.9 Mbp), RTD (5.9-8.2 Mbp), and core (1.9-5.9 Mbp) regions. Rows 5-8: line plots of significant and non-significant upstream (green) and downstream (purple) frontier index values for WT, Δ*lsr2*, Δ*lsrL* and Δ*lsr2*Δ*lsrL* strains. Vertical black dashed lines indicate constitutive frontier indices conserved across genotypes. The x-axis in all plots denotes genomic coordinates binned at 30 kb resolution.

To quantitatively define the domain boundaries observed in the contact maps, we next examined the formation of chromosomal interaction domains (CIDs). Using an approach adapted from Lioy *et al*. (2021) [27], we computed a ‘frontier index’ to assess whether a given locus contributes to boundary formation, potentially influenced by Lsr2 and/or LsrL (**Figure 5B**). To enable comparisons across genotypes, frontier indices were calculated for each strain and evaluated based on whether upstream (green) and/or downstream (purple) significantly deviated from the WT significance threshold (see Methods for details). In total, we identified 13 CID boundaries in the WT strain, with a majority (8/13) located in the core region of the chromosome. Many of these boundaries were located near ribosomal operons, often falling in between them, rather than directly overlapping with the bins encompassing the operons (**Figure 5B, 6A**). This likely reflects a limitation of the lower resolution (30 kb) of our contact maps. The remaining CID boundaries included two in the LTD, and three in the RTD. Notably, one boundary was detected at ∼6 Mbp – corresponding to the RTD boundary – while another was found at ∼1 Mbp within the LTD. Interestingly, the boundary near 1 Mbp demarcated the transition previously observed in **Figure 2A** where a subregion within the LTD (from ∼1-1.9 Mbp) displayed a protein occupancy profile that was more similar to the core region than the LTD (see **Figure 2A** and **5B** above). This suggests the existence of a structurally distinct subregion within the LTD.

**Figure 6.**
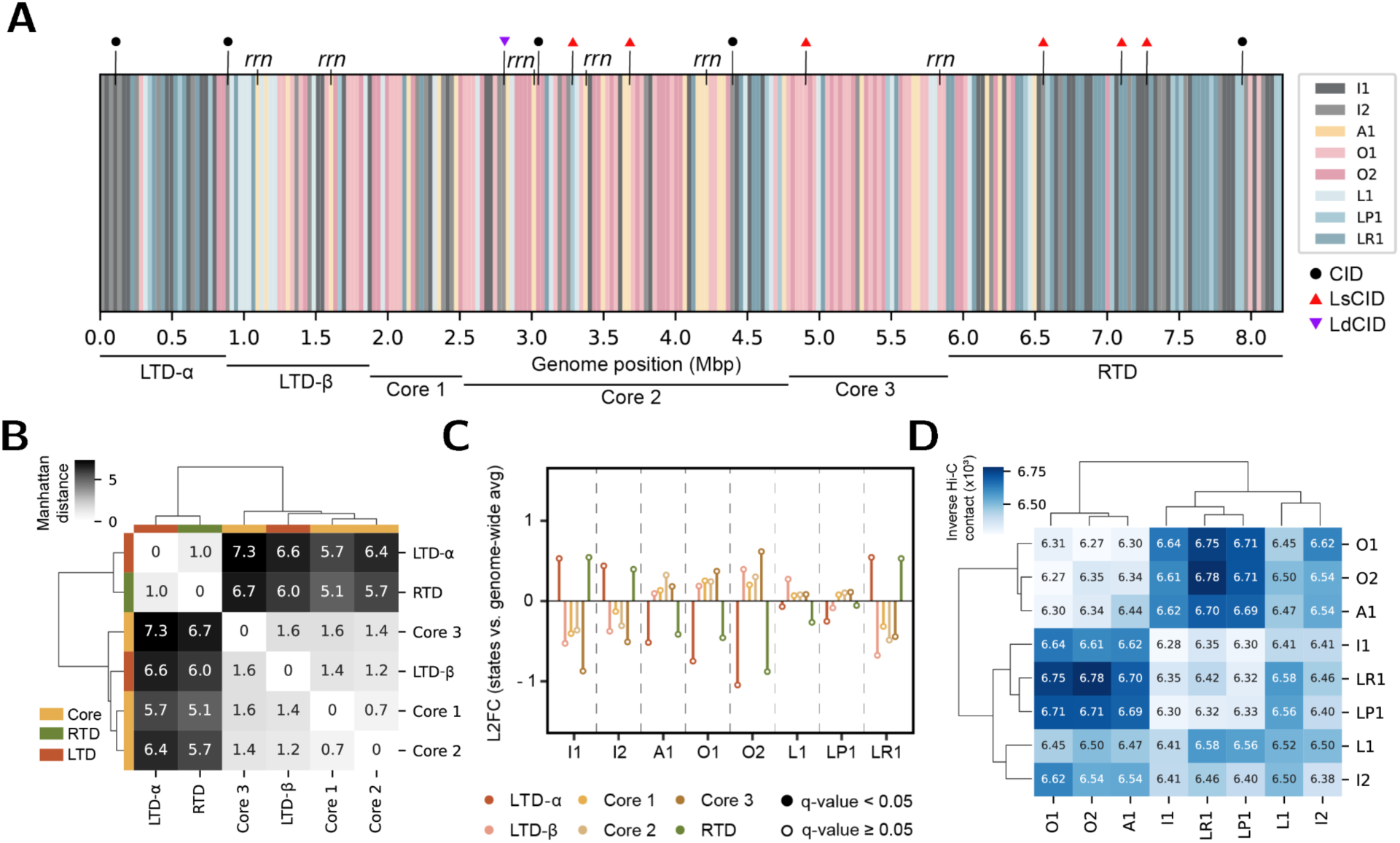
Chromatin state profiles reveal regional subdomains. (**A**) Chromatin states inferred from RNApol ChIP-seq, IPOD-HR, and RNA-seq data are shown at 30 kb resolution across the chromosome, depicted as the most enriched state per bin. Labeled tracks include ribosomal operons, large-scale chromosomal regions (LTD, RTD, and core region), and subregions (LTD-α, LTD-β, Core 1-3). Subregions are defined as: LTD-𝛼: 0-0.9 Mbp; LTD-𝛽: 0.9 - 1.9 Mbp; Core-1: 1.9 - 2.5 Mbp; Core-2: 2.5 - 4.8 Mbp; Core-3: 4.8 - 5.9 Mbp; RTD: 5.9 - 8.2 Mbp. CID boundaries identified for WT in Figure 5B are labeled as either constitutive CID boundaries (CID; black) representing CID boundaries identified in WT, Δ*lsr2*, Δ*lsrL*, and Δ*lsr2*Δ*lsrL* strains; Lsr2-L suppressed CID boundaries (LsCID; red) representing CID boundaries that appeared in all mutants (Δ*lsr2*, Δ*lsrL*, and Δ*lsr2*Δ*lsrL*) only; and Lsr2-LsrL-dependent CID boundaries that disappeared in all mutants (LdCID, gray). (**B**) Heatmap showing pairwise similarities in chromatin state composition across subregions based on Manhattan distances calculated from the proportion of 5 bp bins assigned to each state. Regions are hierarchically clustered based on Manhattan distance and colored by domain type: LTD (red), RTD (green), and core region (orange). (**C**) L2FC of the proportion of chromosomal subregions within each assigned chromatin state relative to the genome-wide average, based on 5 bp resolution bin assignments. Lollipop heads indicate statistical significance from an approximate permutation test (10,000 iterations; filled: q-value < 0.05, unfilled: q-value ≥ 0.05). (**D**) Heatmap of median spatial distances between chromatin state pairs inferred from WT Hi-C contact matrix at a 30 kb bin resolution. Distances are computed as the log10-transformed inverse Hi-C contact (×10^3^) between bins assigned to each chromatin state, excluding local interactions (± 750 kb). Lower inverse Hi- C contacts values indicate closer spatial proximity (or, more properly, lower frequency of close contact between loci), while higher values reflect greater spatial proximity between chromatin states. States are clustered by Manhattan distance to reveal 3D spatial compartmentalization.

The loss of *lsr2* alone resulted in the appearance of 27 new CID boundaries not observed in the WT strain, including 12 in the core region, 6 in the LTD, and 9 in the RTD (**Figure 5B**). Many of these newly formed CIDs in the Δ*lsr2* mutant localized near BGCs, which was consistent with a role for Lsr2 in compacting and repressing these loci. Loss of Lsr2 likely led BGCs to form distinct structural domains as a result of their derepression and increased transcriptional activity (as observed in **Figures 2-4**). In comparison, we identified 14 new CID boundaries in the Δ*lsrL* mutant (1 in the LTD, 8 in the core, 5 in the RTD), many of which were a subset of those observed in the Δ*lsr2* mutant. The Δ*lsr2*Δ*lsrL* mutant exhibited 21 new boundaries (4 in the LTD, 10 in the core, 7 in the RTD), displaying a similar profile to that of the Δ*lsr2* mutant. Across all genotypes, we identified five constitutive CID boundaries that were consistently present in WT and all mutant strains, indicating their independence of Lsr2 and LsrL functions (**Figure 5B, 6A**). In contrast, six CID boundaries, located in the core and RTD regions, were shared across all three mutants but were absent in WT, identifying them as Lsr2/LsrL suppressed CIDs (LsCIDs; **Figure 5B, 6A**). Their consistent emergence in all of the mutant strains implies that Lsr2 and LsrL normally act together to prevent boundary formation at these sites. Finally, we identified a single CID boundary near ∼2.5 Mbp that was lost across all mutant strains, which we defined as an Lsr2/LsrL dependent CID (LdCID), as Lsr2 and LsrL are both required to maintain this domain structure (**Figure 5B, 6A**).

Overall, the results obtained here on the chromosomal structure in the presence of *lsr2* and/or *lsrL* deletions reinforced our earlier observations at the chromatin level: Lsr2 has a predominant role in shaping chromosomal structure and gene expression both locally in BGCs and globally, while LsrL’s effects appeared dependent on Lsr2 presence; we also identified several CID boundaries that depended on Lsr2 and LsrL both being present. Moreover, the observation that the Δ*lsrL* mutant shared many CID boundaries with the Δ*lsr2* mutant further supported our model in which Lsr2 compensates for the loss of LsrL at various loci.

### Chromatin state patterns uncover subregional organization within the core and peripheral regions

Building on the observation of a structural transition around ∼1 Mbp within the LTD (**Figure 2A, 5B**), we next investigated whether similar substructuring existed in the core and RTD. To explore this, and to directly compare chromatin states with Hi-C contact frequencies, we conformed the resolution of the chromatin state assignments to match our Hi-C data by aggregating state assignments into 30 kb bins, assigning each window the most enriched chromatin state (**Figure 6A**). This approach allowed for the visualization of large-scale structural trends across the chromosome and characterization of region-level patterns.

Based on the chromatin state distribution along the chromosome, we observed that the region spanning 0-0.9 Mbp displayed a chromatin landscape distinct from that of the adjacent 1-1.9 Mbp region (**Figure 6A**). As a result, we propose subdivision of the *S. venezuelae* LTD into two subregions: LTD-𝛼 and LTD-𝛽. LTD-𝛼 exhibited a chromatin state profile more similar to that of the RTD, with both enriched for Lsr2-regulated chromatin states such as L1, LR1, and LP1. In contrast, LTD-𝛽 more closely resembled the chromatin state landscape of the core region. Within the core, we observed non-uniform chromatin state patterns that motivated subdivision into three manually defined subregions. Core-2 was readily distinguishable due to its high density of ribosomal operons and enrichment for transcriptionally active states (A1, O1, O2). Notably, Core-2 was flanked by regions enriched for inactive and/or Lsr2-regulated states (I1, I2, L1, LP1 and LR1), which overlapped with BGCs encoded in those regions (**Figure 5B**). In contrast, core-1 exhibited a more heterogeneous profile of open chromatin, Lsr2-regulated and inactive states, whereas core-3 was dominated by open chromatin states with only localized Lsr2-regulated states near several BGCs. Unlike the LTD, the RTD did not exhibit evidence of subregional chromatin patterning, and thus remained as a large terminal region with the previously defined RTD boundaries.

To further evaluate the chromatin state landscape within the subregions that we identified, we quantified chromatin state composition at the original 5 bp resolution using pairwise Manhattan distances (**Figure 6B**). These results further confirmed the distinctiveness of each region: the terminal regions LTD-𝛼 and RTD clustered together, reflecting similar chromatin state profiles, while core-1, core-2, and core-3 formed a separate cluster, together with LTD-𝛽. Although statistical significance was not reached in chromatin state enrichment analyses (**Figure 6C**), again likely due to the 5 bp resolution of states relative to the large domain sizes, we observed clear trends: core subregions were enriched for highly transcribed states (O1, O2) and depleted for transcriptionally inactive (I1, I2) or Lsr2-regulated states while LR1 remained highly enriched in LTD-𝛼 and RTD. The emergence of subregional chromatin domains indicates that within large domains, internal structures can vary significantly, particularly in domains like LTD-𝛼 and RTD where Lsr2-associated chromatin states were the most enriched, underscoring the importance of Lsr2 in shaping terminal chromosomal architecture.

Given that Lsr2 can mediate its regulatory effects via bridging adjacent loci [31], we integrated the chromatin state assignments (at 30 kb resolution) with the WT Hi-C contact matrix to investigate whether transcriptionally active or Lsr2-regulated regions preferentially associated in 3D space, beyond local contacts (± 750 kb; **Figure 6D**). We found that chromatin states with similar transcriptional profiles, such as active states (O1, O2, A1) or inactive/Lsr2-associated states (I1, I2, L1, LP1, LR1), tended to interact more frequently over short genomic distances – *i.e.*, we observed spatial clustering in three-dimensional space of regions with similar chromatin states. In contrast, interactions between dissimilar chromatin states, such as active versus Lsr2-regulated regions, were generally more separated, reflecting reduced spatial proximity between transcriptionally distinct domains. Comparable spatial organizational features of chromatin state clustering were observed in the Δ*lsr2*, Δ*lsrL*, and Δ*lsr2*Δ*lsrL* Hi-C contact matrices (**Supplementary Figure 5A-C**). Overall, this pattern was consistent with spatial compartmentalization, where chromosomal regions with similar transcriptional or regulatory domains are preferentially clustered in 3D space for efficient functioning [43–46]. The clustering of similar states likely reflects local chromatin environments that promote coordinated regulation.

## Discussion

Lsr2 has recently emerged as a key regulator of specialized metabolic genes in *Streptomyces*, playing a dual role in genome compaction and transcriptional repression through its ability to bind and oligomerize and/or bridge DNA segments, thereby both locally repressing transcription and altering chromosomal topology [31]. Despite this growing understanding of Lsr2 functions, how Lsr2 impacts the local and global architecture of the chromosome remains unknown, and the role of its highly conserved paralog, LsrL, has been largely unexplored.

In this study, we reinforce the established role of Lsr2 as a global and locus-specific repressor, silencing genes within and beyond BGCs [32]. We extend this understanding by showing that its repressive effects are especially pronounced in the core and left chromosomal arm, indicating more spatial bias in its regulatory activity than previously recognized. In contrast, LsrL appears to have a more modulatory regulatory function as it influenced a much smaller subset of genes; most of these were also targets of Lsr2. Many of the genes that were downregulated in the Δ*lsrL* mutant were located in the RTD, a pattern that was not observed for either the Δ*lsr2* single or the Δ*lsr2*Δ*lsrL* double mutant. These findings indicate that the regulatory impacts of LsrL activity are not simply redundant with those of Lsr2, but that it may have locus-dependent activity and in some cases antagonistic effects, promoting transcription at genes in specific regions of the chromosome where Lsr2 exerts repression. In other cases, however, we observed that LsrL acted in a supplementary capacity, contributing to repression alongside Lsr2 (*e.g.*, the chloramphenicol biosynthesis cluster).

Previous studies have shown that the loss of *lsr2* in wild *Streptomyces* isolates leads to the activation of numerous metabolic pathways [31,33]. These observations, and our data here, suggest that Lsr2 is likely the primary repressor of many genomic regions encoding genes involved in specialized metabolism, and that additional transcriptional activators, such as CmIR in the chloramphenicol cluster, can counter-silence the repressive effects of Lsr2 by recruiting RNApol to nearby promoter regions [31]. Our findings here raise the possibility that LsrL may also be subject to counter-silencing at loci where it functions as a co-repressor, and for sites where it acts antagonistically, it could itself contribute to the counter-silencing of Lsr2’s repressive activity.

Our findings consistently suggested that LsrL activity was largely dependent on the presence of Lsr2. The Δ*lsr2*Δ*lsrL* double mutant displayed transcriptional and occupancy profiles nearly identical to those of the Δ*lsr2* mutant, implying that without Lsr2, LsrL’s modulatory capacity is lost. In the Δ*lsrL* mutant, however, protein occupancy at some loci remained higher than in the Δ*lsr2* mutant, which is consistent with the ability of Lsr2 to retain its binding in the absence of LsrL. This implies an asymmetrical regulatory relationship between these two proteins, where Lsr2 can partially compensate for the loss of LsrL, but not vice versa. This co-regulatory relationship is similar to that of the recently described interplay between the nucleoid associated proteins H-NS and TsrA in *Vibrio cholerae*. In this system, TsrA primarily targets virulence-associated genes in concert with H-NS, and deletion of both *hns* and *tsrA* yields a transcriptional profile similar to that of the *hns* single mutant [47]. Unlike H-NS and TsrA, however, we found no evidence for additive effects of regulation at any locus in the Δ*lsr2*Δ*lsrL* mutant. Some of the regulatory effects observed here, particularly those associated with LsrL, may instead arise indirectly through changes in chromosomal organization or transcriptional networks influenced by Lsr2. Future work will focus on dissecting the mechanistic basis underlying the interaction between Lsr2 and LsrL, including simultaneous measurement of Lsr2 and LsrL binding via ChIP-seq to determine whether they physically associate at shared sites, or influence each other’s binding through changes in local DNA topology, and whether they form hetero-oligomeric complexes or remain in homomeric states when bound to DNA.

By integrating chromosome conformation, protein occupancy and gene expression data, we have been able to assign regions of the *S. venezuelae* chromosome to different chromatin states, corresponding to distinct genomic features such as protein-coding genes (both BGCs and non-BGCs), intergenic regions containing TFBSs and promoter regions, and AT-rich regions. Importantly, our chromatin state data was entirely consistent with our Hi-C contact maps, where loss of *lsr2* had a more pronounced effect on short-range contacts. This pattern parallels observations in *Escherichia coli*, where deletion of H-NS similarly increased local contact frequencies, consistent with the loss of bridging interactions that normally link distant loci [1]. In contrast, deleting *lsrL* had a less pronounced effect on short range contacts overall, as would be expected if Lsr2 continues to bridge loci in the absence of LsrL. The increase in short range contacts observed upon loss of Lsr2 may therefore reflect the collapse of higher-order chromosomal loops that it normally maintains. By bridging distant regions, Lsr2 likely contributes to spatial dispersion of DNA loops and overall nucleoid organization; in its absence, these regions compact into localized clusters that appear as enhanced short-range interactions. This reorganization reflects a direct structural role of Lsr2 in maintaining long-range chromosomal organization.

One of the notable changes in chromosome architecture in the Δ*lsr2* mutant was the appearance of multiple new CID boundaries that were associated with (overlapping and nearby) BGCs. The occurrence of these new boundaries is entirely aligned with previous work in *S. ambofaciens* that showed highly transcribed genes can act as boundary elements that segment the chromosome into domains [27]. In *S. venezuelae*, the boundary-forming potential of BGCs appears to be normally masked by Lsr2, which represses BGC transcription and likely maintains compaction at those sites. Loss of *lsr2* lifts this repression, allowing increased transcription and localized decompaction of the chromosome. This in turn appears to enable the formation of new structural boundaries, particularly in regions of the chromosome where BGC density is high.

More broadly, the tendency of chromatin states with similar transcriptional profiles – whether active, inactive, or Lsr2-associated – to cluster in three-dimensional space, while dissimilar states are more separated, suggests that the *S. venezuelae* chromosome exhibits transcription-linked spatial organization. A similar relationship between transcriptional activity and spatial proximity has been observed in *E.coli*, where actively transcribed loci form local 3D domains and RNApol foci, while adjacent H-NS-silenced regions segregate separately [43,48,49]. Within this framework, Lsr2 helps coordinate local domain structure and reinforces higher-order spatial organization at the loci it regulates.

Overall, our study provides substantial insights into the distinct and overlapping regulatory roles of Lsr2 and LsrL in governing genes including specialized metabolite-producing genes within *S. venezuelae*. Understanding the mechanisms governing BGC regulation is a crucial step towards unlocking the full metabolic potential of these chemically gifted bacteria.

Our findings have further shed light on how nucleoid associated proteins work in concert to shape chromosome architecture, mediate nucleoid compaction, and influence the 3D spatial organization of the genome. By linking transcriptional regulation to chromatin state dynamics and higher-order chromosome structure, our findings highlight the multifaceted strategies by which *Streptomyces* coordinate specialized metabolism with global genome organization.

Nevertheless, several limitations and opportunities for future work must be kept in mind. Our study covers a single growth condition, and is limited (as with all applications of the IPOD-HR method) by the fact that we cannot definitively assign observed protein occupancy to any particular protein, and that some proteins which do not crosslink effectively to DNA may be missed [37]. Future follow-ups on the work presented here will likely include both a more detailed dissection of the precise binding sites of Lsr2 and LsrL under different conditions, and expansion to consideration of other *Streptomyces* NAPs such as HupA and/or HupS [50] and their interplay with the Lsr2/LsrL silencing system.

## Materials and Methods

### Bacterial strains and culture conditions

The *S. venezuelae* NRRL B-65442 strain served as the WT and base strain for all experiments conducted in this study. *S. venezuelae* strains were cultured at 30°C on MYM (maltose, yeast extract, malt extract) agar or in MYM liquid media.

### Mutant strain construction

The previous construction of in-frame deletions of *lsr2* (vnz_15890/vnz_RS16005) and *lsrL* (vnz_18875/vnz_RS19015) is described in Gehrke *et al*. (2019) [32]. The Δ*lsr2*/Δ*lsrL* mutant strain was constructed by replacing *lsrL* with an aparamycin resistance cassette in the scarred Δ*lsr2* background [32].

### Sample preparation for IPOD-HR, RNApol ChIP-seq and RNA seq

Spores of each strain (WT, Δ*lsr2*, Δ*lsrL*, and Δ*lsr2*Δ*lsrL*) were inoculated in 20 mL of MYM liquid medium and grown overnight with shaking at 30°C. Subsequently, each strain was subcultured into six independent flasks containing 100 mL of liquid MYM to a final OD_600_ of 0.1 and grown with shaking at 30°C. After 16 h, three 15 mL aliquots were removed from each culture, centrifuged at 7,800×g at 4°C for 5 minutes, and the supernatant discarded. These aliquots were stored at -80°C and subsequently used for RNA extraction (see below). One of the six replicates served as the test sample for determining optimal DNase treatment conditions for the IPOD-HR and RNApol ChIP-seq experiments. For the remaining five replicates, the three replicates that yielded a complete set of the highest quality DNA and RNA samples for IPOD-HR, RNApol ChIP-seq, and RNA-seq were sent for next-generation sequencing (described below). This approach ensured that data were collected from the same cultures across each experiment.

To the remaining 50 mL of culture, rifampicin was added to a final concentration of 500 μg/mL and the cultures were incubated with shaking at room temperature for 10 min (rifampicin was used to trap RNA polymerase at the promoter and prevent further transcription initiation).

Formaldehyde was then added to a final concentration of 1% (w/v) to induce protein and DNA crosslinking, and the cultures were incubated for an additional 30 min at room temperature.

Next, glycine was added to a final concentration of 125 mM to quench the crosslinking, after which the cultures were incubated for another 5 min. The cultures were then centrifuged in 50 mL Falcon tubes at 7,800×g at 4°C for 5 min, after which the supernatant was discarded. The resulting cell pellets were washed three times with phosphate-buffered saline (PBS), by resuspending the pellets in 20-40 mL cold PBS and centrifuging as above.

### Cell lysis and DNA preparation for IPOD-HR and RNA polymerase chromatin immunoprecipitation

We performed the IPOD-HR interface extraction and nucleic acid purification, RNA polymerase chromatin immunoprecipitation, and DNA extraction following reverse crosslinking, as detailed in Freddolino *et al*. (2021) [37], with modifications to the protocol detailed below.

#### Lysis of S. venezuelae cell pellets

Each of the washed cell pellets from above were resuspended in 1.5 mL lysis buffer (10 mM Tris-HCl pH 8.0, 50 mM NaCl, 14 mg/ml lysozyme, 0.8% Triton X-100, 1× protease inhibitor) and incubated at 37°C for 30 min. The samples were incubated on ice for 2 min, then sonicated for six cycles of 15 s continuous sonication followed by 45 s on ice (output 40%, 4). Cell debris was removed by centrifugation for 20 min at 16,000×g at 4°C and the supernatant was transferred to a 15 mL Falcon tube (∼1.8 mL).

#### Determination of optimal DNase treatment conditions

One of the six replicates was used to determine the appropriate conditions for DNase treatment that would yield DNA fragments between 100-200 bp, as we found that required treatment was different for each sample. To determine the appropriate time for DNase treatment, 200 µL of 10× Turbo DNase Buffer, 10 µL Turbo DNase, and RNase A to a final concentration of 15 µg/mL were added to this sample, and incubated for 90 min at 37°C. Every 10 min during the 90-min time course, a 50 µL aliquot was collected from each sample, mixed with 10 µL of 0.5 M EDTA (pH 8), and incubated on ice to inhibit further DNase digestion. Once all aliquots had been collected, they were subjected to phenol-chloroform extraction, and 5-10 µL of the upper phase was run on a 2% agarose gel. The time that yielded the most DNA between 100-200 bp was used to digest the remaining five replicates.

#### DNase treatment

The remaining five replicates were DNase-treated using the experimentally determined optimal conditions. After DNase digestion, 500 µL of 0.5 M EDTA (pH 8.0) was added to quench the reaction. The samples were clarified by centrifuging at 7,800×g for 10 min at 4°C. A 50 µL aliquot (IPOD input DNA) was removed and diluted 1:9 in elution buffer (50 mM Tris, pH 8.0; 10 mM EDTA; 1% SDS) and kept on ice until the reverse crosslinking step. A 100 µL aliquot of the clarified lysate was phenol-chloroform extracted and 5 µL of the upper (DNA) phase was run on a 2% agarose gel to confirm that the DNase treatment was successful and had yielded DNA with a maxima centered between 100-200 bp.

#### IPOD-HR

One milliliter of the resulting clarified lysate was transferred to a 15 mL Falcon tube and mixed with an equal volume of 100 mM Tris base. An equal volume of 25:24:1 phenol:chloroform:isoamyl alcohol was added, the sample vortexed, and incubated for 10 min at room temperature. After incubation, the sample was centrifuged at 7,800×g for 5 min at 4°C, allowing the formation of a white disc at the aqueous–organic interface enriched for protein–DNA complexes.

Both the upper and lower aqueous phases were removed with a pipette and discarded. The remaining interface disc was washed with 350 µL TE buffer (10 mM Tris, pH 8.0; 1 mM EDTA), 350 µL 100 mM Tris base, and 700 µL 24:1 chloroform:isoamyl alcohol. The sample was vortexed and centrifuged as above. The upper and lower aqueous phases were removed again, and the interface disc was washed with 1 mL TE buffer and 1 mL 24:1 chloroform:isoamyl alcohol. The sample was vortexed and centrifuged as above, and both upper and lower aqueous phases were discarded. The interface disc was resuspended in 700 µL of elution buffer by vortexing. To reverse protein-DNA crosslinking, the input and IPOD-HR samples were incubated at 65°C overnight.

The next day, the samples were allowed to cool, and RNase A was added to each input and IPOD-HR sample to a final concentration of 60 µg/mL. The samples were incubated at 37°C for 2 h. Proteinase K was then added to a final concentration of 120 µg/mL and incubated at 50°C for 2 h. DNA was phenol-chloroform extracted, the upper phase was transferred to a clean tube and precipitated by the addition of 2 volumes of cold 95% ethanol and 0.1 volume of precipitation buffer (5 M CH_3_COONa/CH_3_COOH, pH 5.2) and incubation at -20°C overnight. The next day, samples were centrifuged at 16,000×g for 30 min at 4°C, and the supernatant discarded. The DNA pellet was washed with ice-cold 70% ethanol and dried by placing the tube up-side-down on a paper towel. The DNA pellet was resuspended in 30 µL 10 mM Tris-HCl, pH 8.0.

#### RNA polymerase chromatin immunoprecipitation

Protein A-Sepharose beads were prepared in 1.5 mL tubes by resuspending 125 mg of the beads in 1.2 mL of Buffer A (0.02 M NaH_2_PO_4_, 150 mM NaCl, pH 8.0) and incubating at room temperature for 30 min. The beads were then aliquoted into separate 1.5 mL tubes for each sample (100 µL slurry per sample). The beads were equilibrated by removing the storage buffer and adding 1 mL of IP buffer with BSA (100 mM Tris pH 8.0, 300 mM NaCl, 2% Triton X-100, 0.1 mg/mL Bovine Serum Albumin (BSA)). Beads were centrifuged at 1,000×g for 1 min at 4°C and allowed to settle on ice for 2 min. This wash step was repeated a second time, and the beads were resuspended in 100 µL IP buffer with BSA.

To pre-clear the lysate, the remaining clarified lysate from after DNase treatment (∼1 mL) was added to the equilibrated beads and incubated with gentle shaking at 4°C for 2 h. The beads were pelleted as above, the supernatant (pre-cleared lysate) was transferred to a clean tube, and 10 µL of anti-RNA polymerase antibody (custom production run equivalent to NeoClone WP0023, clone 8RB13) was added to each sample. The samples were incubated overnight with gentle shaking at 4°C. After removing the lysate, the beads were washed three times with 1 mL of Buffer A and stored at 4°C.

The next day, the beads were equilibrated as before in IP buffer with BSA. The antibody-lysate mixture was added to the beads and incubated with gentle shaking at 4°C for 4 h. The beads were centrifuged as above and the majority of the supernatant discarded by removing with a pipette. The beads were washed at room temperature with 1 mL of each of the following solutions in order:

- Wash buffer A (100 mM Tris, pH 8.0; 250 mM LiCl; 2% Triton X-100; 1 mM EDTA)
- Wash buffer B (100 mM Tris, pH 8.0; 500 mM NaCl; 1% Triton X-100; 0.1% sodium deoxycholate; 1 mM EDTA)
- Wash buffer C (10 mM Tris, pH 8.0; 500 mM NaCl; 1% Triton X-100; 1 mM EDTA)
- IP buffer with EDTA (100 mM Tris, pH 8.0; 300 mM NaCl; 2% Triton X-100; 1 mM EDTA)
- TE buffer (10 mM Tris, pH 8.0; 1 mM EDTA)

After washing, the beads were resuspended in 500 µL elution buffer and protein-DNA crosslinking was reversed by incubating the samples at 65°C overnight. Samples were treated with RNase A and proteinase K, and the DNA precipitated as above for the input and IPOD-HR samples.

### Illumina library preparation for IPOD-HR and RNA polymerase and chromatin immunoprecipitation

All purified DNA samples were run on an Agilent TapeStation to assess DNA quality prior to library preparation. Libraries were prepared for Illumina sequencing using the NEBNext Ultra II DNA kit (New England Biolabs). The samples were pooled into libraries and sequenced on an Illumina NextSeq 2000, in a P3, 2×50bp configuration at the McMaster Genomics Facility.

### IPOD-HR data analysis pipeline

#### Read pre-processing

Raw NGS reads were processed using a combination of read quality filtering, adapter trimming, alignment and mapping, and protein occupancy using the IPOD-HR pipeline (version 2.8.1) via a singularity container (accessible at https://github.com/freddolino-lab/ipod) [37]. Briefly, sequencing adapters were removed using Cutadapt [51] (version 4.0), poor quality reads were trimmed with Trimmomatic [52] (version 0.39) using the parameters “TRAILING:3 SLIDINGWINDOW:4:15 MINLEN:10”.

#### Alignment and protein occupancy calling

Remaining reads were aligned to the *S. venezuelae* NRRL B-65442 NZ_CP018074.1 reference genome using bowtie2 [53] (version 2.4.4) with “very sensitive, end to end” presets and dovetail alignments allowed. Only concordant pairs were used for downstream quantitation. The surviving processed reads were assessed using FASTQC [54] for sequence content, quality, and duplication and bowtie2 outputs. The quality of alignments were evaluated using bowtie2 output metrics, including the overall alignment rates. Read densities were quantile normalized, and the log ratios of IPOD-HR and RNApol ChIP-seq were calculated at a 5 bp resolution relative to the corresponding input sample. To isolate occupancy scores attributable to DNA-binding proteins distinct from RNApol, the IPOD-HR pipeline subtracts RNApol ChIP-seq normalized robust z-scores (RNApol ChIP-seq vs. input) from the IPOD normalized robust z-scores. The resulting bedgraph files, containing enrichment scores for RNApol ChIP-seq and IPOD-HR, were used for downstream analyses. To visualize occupancy changes for both RNApol ChIP-seq and IPOD-HR across the *S. venezuelae* chromosome in Figure 2A and Supplementary Figure 1A, 20-kb rolling averages were calculated from the 5-bp resolution occupancy score bedgraph files using the window and map commands in bedtools (version 2.30.0) [55].

### RNA isolation, rRNA depletion, and RNA-sequencing library preparation

Using the cell pellets that were set aside from the cultures used for IPOD-HR and RNApol ChIP-seq, RNA was extracted from three to five replicates each of WT, Δ*lsr2*, Δ*lsrL*, and Δ*lsr2*/Δ*lsrL* mutant strains. RNA was isolated as described in Moody *et al*. (2013) [56], using a modified guanidium thiocyanate protocol [57]. Primers amplifying a 144 bp region of *rpoB* (*vnz_21470*) were used for PCR checks, alongside a quantified chromosomal DNA control, to confirm the absence of DNA contamination. RNA quality was assessed first by agarose gel electrophoresis, and subsequently using an Agilent 2100 Bioanalyzer. In the case where RNA was successfully extracted from more than three replicates, those with the highest quality were used for sequencing.

Purified RNA was depleted of rRNA using the Bacterial NEBNext rRNA Depletion Kit (New England Biolabs). Library preparation was conducted using the NEBNext Ultra II Directional RNA kit (New England Biolabs). The samples were pooled into libraries and were sequenced on an Illumina NextSeq 2000, P3, in a 2×50bp configuration at the McMaster Genomics Facility.

### Lsr2 ChIP-seq data processing

Previously published Lsr2 ChIP-seq data from *S. venezuelae* were downloaded from the NCBI Gene Expression Omnibus (GEO accession: GSE115439) [32]. Two biological replicates of the FLAG-tagged Lsr2 ChIP-seq strain (Δ*lsr2* + Lsr2-3×FLAG) and two replicates of the untagged Δ*lsr2* control strain were processed using the same pipeline described for IPOD-HR and RNApol ChIP-seq analysis. Briefly, sequencing reads were adapter- and quality-trimmed using Cutadapt [51] (version 4.0) and Trimmomatic [52] (version 0.39), then aligned to the *S. venezuelae* NRRL B-65442 reference genome (GenBank accession: NZ_CP018074.1) using bowtie2 [53] (version 2.4.4) with “very sensitive, end-to-end” alignment settings. Only concordantly mapped read pairs were retained. For each replicate, genome-wide read coverage was computed at a 5 bp resolution using the window and map commands of bedtools (version 2.30.0) [55]. Coverage values were log2-transformed and mean centered, and biological replicates were averaged separately for each sample. The resulting occupancy profiles were used solely for visualization purposes and were excluded from downstream analyses.

### RNA-sequencing analysis

#### Read pre-processing and quality control

The raw RNA-sequencing reads were processed using the IPOD-HR pipeline, beginning with the removal of Illumina adapter sequences with cutadapt [51] (version 4.0) and trimming of low quality reads with trimmomatic [52] (version 0.39), as described above. High quality reads were evaluated using FASTQC [54] for quality, sequence duplication, and sequence content.

#### Transcript abundance quantification and differential expression analysis

Transcript abundance from processed RNA-seq reads was estimated in transcripts per million (TPM) units using Kallisto [58] (version 0.44.0). Although rRNA depletion was performed experimentally, rRNA and tRNA gene annotations were also removed from the gene-level annotation FASTA file of the *S. venezuelae* NRRL B-65442 NZ_CP018074.1 reference genome to further ensure that transcript abundance estimates were not biased by residual rRNA or tRNA. Additionally, each gene sequence was extended by 50 bp upstream and downstream in a strand-aware manner to account for potential reads overlapping gene boundaries. The modified annotation fasta file was then used to construct the Kallisto index with a kmer size of 17 (“-k 17”). RNA-seq reads were pseudoaligned and quantified with Kallisto [58] (version 0.44.0) with default settings save for a change in number of bootstraps: “-b 100, -rf-stranded”. The quality of pseudoalignment was evaluated across all samples using Kallisto metric outputs, including percent pseudoaligned and percent unique. To assess replicate-to-replicate consistency, Spearman’s rank correlation coefficients were calculated in Python 3.10.10 based on the quantified transcript abundances. Due to the inconsistency and poor quality of replicate 3 in the Δ*lsrL* mutant, based on Spearman’s rank correlation coefficient, and pseudoalignment metrics, replicate 3 was removed, leaving three replicates for WT, three replicates for Δ*lsr2*, two replicates for Δ*lsrL*, and three replicates for Δ*lsr2*Δ*lsrL* for gene expression analysis.

Differential gene expression calling of quantified reads was performed using the sleuth [59] (version 0.30.1) R package, employing the Sleuth response error measurement (full) model with the formula “∼genotype” where *genotype* represents the four conditions (WT, Δ*lsr2*, Δ*lsrL*, and Δ*lsr2*Δ*lsrL*). The Wald test was used to quantify differential expression values between each genotype and the WT. Principal component analysis (PCA) was performed to explore patterns of gene expression and assess similarities both within and between genotypes. Downstream analyses were performed with in-house R 4.2.0 and Python 3.10.10 scripts. To determine the gene targets similarly regulated between Lsr2 and LsrL proteins, the Spearman rank correlation coefficient within Python 3.10.10 was performed on gene expression profiles between Δ*lsr2*, Δ*lsr2*, and Δ*lsr2*Δ*lsr2* mutants.

#### RNA read coverage

To generate high-resolution occupancy plots, reads were instead aligned with bowtie2 to the *S. venezuelae* NRRL B-65442 NZ_CP018074.1 reference genome as noted above for DNA reads. The subsequent read occupancies were calculated using the bedtools [55] 2.30.0 genomecov command. To account for differences in sequencing depth across the replicates across all samples, we normalized the quantified read coverages using a Trimmed Mean of the M-values (TMM) approach [60]. We first calculated the size factors (effect size estimation) by removing the top and bottom 5% of coverage values and calculating the mean of the remaining coverage values. To apply normalization in each replicate, we divided the coverage values by the corresponding size factor, ensuring that differences in sequencing depth would not bias downstream analyses. The normalized coverage value was log2 transformed using a pseudocount of 0.01 to avoid taking the log of zero. The resulting normalized coverage values were utilized for downstream analyses, including the generation of occupancy plots.

### GO term enrichment analysis with iPAGE

To perform the GO term enrichment analysis based on L2FCs for the Δ*lsr2* and Δ*lsrL* mutants relative to WT, we extracted GO terms from the *S. venezuelae* NRRL B-65442 NZ_CP018074.1 Genbank file using a custom Python script. This resulted in 912 unique GO terms across 1,685 genes. Enrichment analysis was conducted using iPAGE [42] (version 1.2c), via a singularity container. The input file consisted of two columns: the gene label and the corresponding Wald test statistic representing the estimated effect size (L2FC; “b” value) generated by Sleuth RNA-seq analysis. Given that the Wald statistic was used as a continuous variable, iPAGE was run in continuous mode with data discretized into five bins (--extype=continuous --ebins=5 --independence=0 --max_p=0.05).

### HMM for chromatin state inference

HMM fits were performed using the hmmlearn 0.3.0 python package with Gaussian emissions. The genome, spanning 8,222,198 bp, was analyzed at a resolution of 5 bp, resulting in a total of 1,644,440 observations per genotype (WT, Δ*lsr2*, Δ*lsrL* and Δ*lsr2*Δ*lsrL*) across the genome for each data type with input features as RNApol ChIP-seq robust z-scores, IPOD-HR robust z-scores, and the log2 TMM-normalized read coverage values. To identify the optimal number of hidden states a 5-fold cross-validation was performed, partitioning the genome into windows of 1,678 bp, chosen to capture binding peaks and allow the model to learn regulatory patterns over a broad range. This windowing resulted in 4,900 windows per genome. These windows were then evenly divided into 5 folds using a round-robin algorithm, with each fold containing 980 windows, equating to 328,888 observations per fold for each data type. Using mean absolute error to evaluate the model performance across the trained HMMs ranging from 2 to 13 states, we found that a 9-state model provided the best fit with minimal error (Supplementary Figure 1B). We therefore selected a 9-state model to provide a balance of interpretability and predictive performance.

After selecting the 9-state model, we fit a final HMM model using the entire genome across all data types and genotypes resulting in a matrix size of [1,644,440 x 12]. The final state assignments were then obtained using the Viterbi algorithm in hmmlearn with default parameters. After assessing the median of normalized enrichment scores, standard deviation of normalized enrichment scores, and the transition state probabilities in the final 9-state model, we observed that two states (0 and 2) oscillated between each other, with adjacent genomic positions frequently alternating between these two states (state 0 <-> state 2 transition probability > 0.8; Supplementary Table 2). This led us to combine these two states into a single state (O2), resulting in a final 8-state HMM model that was displayed in Figure 2.

### Genomic feature analyses

Genomic features from the *S. venezuelae* genome were analyzed to examine their distribution within chromatin states inferred from the Gaussian HMM. The relative proportion fold change (RPFC) for each genomic feature across chromatin states was calculated relative to the genome-wide proportion of each chromatin state using the following equation:

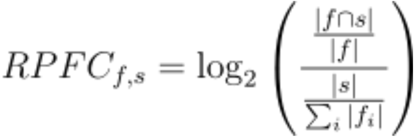

Where *f* denotes the genomic feature (*e.g.*, gene bodies, intergenic regions, promoter regions) and *s* denotes the chromatin state assigned to the region. |𝑓 ∩ 𝑠| denotes the number of regions assigned to chromatin state *s* within genomic feature *f*. |𝑓| denotes the total number of regions for genomic feature *f*. |𝑠| denotes the total number of regions assigned to chromatin state *s* across all genomic features. 𝛴_i_|𝑓_i_| denotes the total number of regions across all genomic features. Each feature was assigned using a combination of approaches as described below.

#### TFBSs

Curated and predicted TFBS sequences and coordinates for *S. venezuelae* were obtained from the LogoMotif database [61], accessible via the public database (https://logomotif.bioinformatics.nl/) containing the ATCC 10712 strain (GCF_000253235.1).

This yielded a total of 3,810 TFBSs across the genome for 82 regulators. To refine these regions, we used bedtools [55] (version 2.30.0) to expand the coordinates by 100 bp upstream and downstream of the TFBS (-100 to +100 relative to TFBS window) and to extract the associated nucleotide sequences. The nucleotide sequences of each TFBS were then mapped from the *S. venezuelae* ATCC 10712 strain to the *S. venezuelae* NRRL B-65442 strain using minimap2 [62] (version 2.28-r1209) with the parameter “-x sr” for short-”read” alignment. This mapping step resulted in ∼94% (3,583) of the TFBSs aligned to the target genome, which were subsequently used for downstream analyses.

#### AT rich regions

AT content was calculated by averaging nucleotide composition across a 21 bp window, with each nucleotide position centered within the window, throughout the *S. venezuelae* genome. To define a threshold boundary of AT rich genes, we fit a Gaussian mixture model of the averaged AT content within the Mclust R package [63] (version 6.1.1). The GMM identified 8 clusters with the most extreme cluster having a boundary of 0.4545. Genomic regions with AT content ≥ 45.45% were thus classified as AT rich regions.

#### Promoter regions

Promoter regions were inferred from predicted operons, which were identified computationally from the *S. venezuelae* NRRL B-65442 genome annotation. Genes located on the same strand and within 200 bp of one another were merged using bedtools [55] (version 2.30.0) to define putative operons. Promoter regions were then defined as 250 bp upstream and 50 bp downstream of the start site of the first gene in each predicted operon, accounting for strand orientation. The corresponding genomic coordinates and nucleotide sequences were extracted for downstream analyses.

#### BGCs

Specialized metabolite BGCs were identified genome-wide using AntiSMASH [38] (accessed via https://antismash.secondarymetabolites.org) with default parameters. This resulted in 32 BGCs associated with 24 types of specialized metabolites. Annotations of data from AntiSMASH were used for subsequent analyses.

#### Chromosomal regions

Coordinates for chromosomal regions, including the LTD, core region, and RTD were obtained from Szafran *et al*., 2021 [7]. These regions were defined for *S. venezuelae* NRRL B-65542 with the following coordinates: LTD (0-1.9 Mbp), core region (1.9-6.3 Mbp), and RTD (6.3-8.2 Mbp). To capture finer-scale variation, we subdivided these regions into LTD-𝛼, LTD-𝛽, core 1, core 2, and core 3 based on distinct chromatin state profiles inferred by the HMM. The RPFC of each chromatin state within these subregions was calculated relative to the genome-wide average. In addition, we quantified dissimilarity among subregions by calculating Manhattan distances between their chromatin state distributions.

### Permutation test for genomic and chromosomal features

An approximate permutation test with 10,000 iterations was performed to identify whether the RPFC calculated for each genomic feature across all states was significantly different from that of the null distribution. The null distribution was generated by rotating the chromatin state assignments across the chromosome. For each permutation, a random integer between 1 and the chromosome length was chosen, and the chromosome length was cyclically shifted by that number of positions. This preserved the overall distribution of chromatin states, while randomizing their positions relative to the genome and chromosomal region features. The RPFC was calculated for each permutation. P-values were calculated using the following equation:

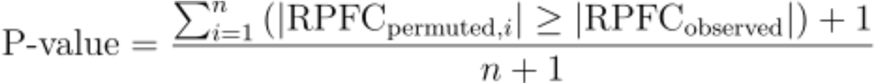

where 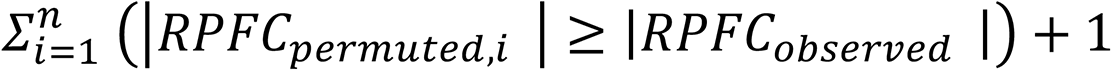 is the total number of permutations that contained RPFCs that were at least as extreme as the original observed RPFC for a given genomic feature across all chromatin states, and n denotes the total number of permutations. To correct for multiple testing, q-values were calculated using the Benjamini-Hochberg method.

For chromosomal regions (LTD, RTD and core region) and subregions (LTD-alpha, LTD-beta, core 1, core 2, and core 3), an alternative permutation approach was used to preserve local autocorrelation in chromatin state assignments. Using the autocorrelation function of the HMM-inferred state sequence, we identified an elbow point corresponding to a correlation length of 8,765 bp, representing the lag at which autocorrelation between chromatin state assignments decays and neighboring regions are no longer strongly correlated. The genome was then segmented into non-overlapping blocks of 8,765 bp, and chromatin states were permuted by cyclically shifting the order of these blocks by a random integer between 1 and the total number of blocks in each iteration. This approach maintained the local dependency structure of the chromatin states while randomizing their genomic locations. For each permutation, the RPFC was recalculated, and p-values were assessed using the equation described above. q-values were then computed using the Benjamini-Hochberg method, as described for genomic features.

### Preparation of Hi-C libraries and analysis

For the preparation of *S. venezuelae* Hi-C contact maps, 5 mL MYM cultures were inoculated with 6×10^3^ colony-forming units (CFU). The cultures were grown for 18 h with shaking (180 rpm) at 30 °C. Cells were then collected and processed according to the protocol described by Szafran *et al*. (2021) [7]. The agarose-purified Hi-C libraries were diluted to a 10 nM concentration, pooled together, and sequenced using Illumina NextSeq500 (2×75 bp; paired reads), as offered by the NGS Services sequencing facility (Plan-les-Ouates, Switzerland). Sequencing data were collected and preprocessed by NGS Services using the software as below: NextSeq Control Software 4.0.2.7, RTA 2.11.3 and bcl2fastq2.17 v. 2.17.1.14. The complete data processing was performed using the Galaxy HiCExplorer web server following the protocol described by Szafran *et al*. (2021) [7].

To define the LTD and RTD boundaries, a principal components analysis was performed on the normalized WT Hi-C contact matrix. Principal component 1 was smoothed using a Gaussian filter and the points at which the smoothed principal component 1 profile intersected the x-axis were used to identify domain boundaries.

### CID boundary detection

To identify CID boundaries, we implemented a frontier-index based method adapted from Lioy *et al*. (2021) [27,64]. To reduce noise, Hi-C contacts were first imputed for bins with zero contact counts using iterative neighbor averaging, where each missing value was replicated by the mean of its non-zero neighbors. The resulting contact maps were then log-transformed with a pseudocount (5th percentile of positive contact values) and smoothed using a Gaussian filter (σ = 0.75 bins; equivalent to ∼22.5 kb at 30 kb resolution). Upstream and downstream frontier indices were calculated as the sum of positive vertical and negative horizontal derivatives, respectively, within a ± 750 kb interaction window. This window size was selected based on a contact decay profile of the WT genotype, where average Hi-C contact frequencies plateaued, reflecting the range of local chromosome interactions. To identify candidate boundaries, local maxima were detected independently in the upstream and downstream frontier index profiles.

Peak detection was performed per genotype by identifying bins where the frontier signal exceeded neighboring bins.

To assess significance, thresholds were calculated using the WT genotype separately for the canonical core and terminal chromosomal regions. For each region, the thresholds were defined as the median of WT peak frontier index values plus 1.8 times the standard deviation, where the standard deviation was approximated using 1.48 times the median absolute deviation to reduce the influence of outliers. The WT-derived thresholds were then applied across all genotypes to ensure comparability. Symmetrical CID boundaries were defined as bins exhibiting both a significant upstream and downstream frontier index peak within ± 2 bins. Significant asymmetrical peaks, occurring in only one direction, were retained and labeled accordingly.

Frontier index peaks that remained in all four genotypes were labeled as constitutive CID boundaries. For these analyses, we used the following replicate Hi-C datasets, which contained the fewest bins with zero contact counts: WT rep1, Δ*lsr2* rep1, Δ*lsrL* rep2, and Δ*lsr2lsrL* rep2.

### Chromatin state pairwise spatial distance analysis

To integrate the HMM-derived chromatin states with 3D chromosome structure, we converted the chromatin states to match the 30,000 bp resolution of the Hi-C data. For each bin, the most enriched chromatin state was assigned based on the genome-wide enrichment relative to their global chromatin state frequencies. To assess spatial relationships between chromatin states, pairwise contact frequencies were calculated between all bin pairs in the WT, Δ*lsr2*, Δ*lsrL*, and Δ*lsr2*Δ*lsrL* Hi-C contact matrices, excluding local interactions (± 750 kb interaction window), as determined from the contact decay curve where average Hi-C contact frequency plateaued.

Each bin pair was annotated with the most enriched chromatin state, and the contact values were transformed by computing the log10 of the inverse Hi-C contact frequency, with a pseudocount of 10^-6^. The log10-transformed inverse Hi-C contacts were further used to represent 3D spatial distance. For each chromatin state pair, all nonlocal pairwise spatial distances were compiled, and the median distance was used to summarize their 3D spatial relationship. Hierarchical clustering of chromatin states was then performed using Manhattan distances computed from the resulting median distances to assess similarities in spatial interaction profiles between states. The same replicate datasets were used as described above.

## Acknowledgements

Author contributions:

Vivian Ramirez (Software [lead, Formal analysis [lead], Methodology [equal], Validation [equal], Visualization [equal], Writing-original draft [lead], Writing-review & editing [equal])

Lauren Tiller (Investigation [equal], Writing-review & editing [equal]) Marcin Jan Szafran (Investigation [equal], Writing-review & editing [equal] Xiafei Zhang (Investigation [equal])

Dagmara Jakimowicz (Writing-review & editing [equal]

Marie Elliot (Conceptualization [equal], Supervision [supporting], Writing-review & editing [equal] Lydia Freddolino (Conceptualization [equal], Software [supporting], Formal analysis [supporting], Methodology [equal], Validation [equal], Supervision [lead], Visualization [equal], (Writing-review & editing [equal])

## Data availability

RNA-seq, IPOD-HR, and RNA polymerase ChIP-seq data generated in this study have been deposited in the NCBI Gene Expression Omnibus under accession code GSE310427. The Hi-C data generated in this study are available in the ArrayExpress database under accession code E-MTAB-16021.

## Code availability

Analysis scripts and custom code used in chromatin state inferences and frontier indices calculations for CID boundary detection used in this work are available at https://github.com/vivianra/Lsr2_LsrL_S_venezuelae_analyses.

