## Supplementary Figures for "LsrL modulates Lsr2-induced chromatin structure to tune biosynthetic gene cluster regulation in *Streptomyces venezuelae*"

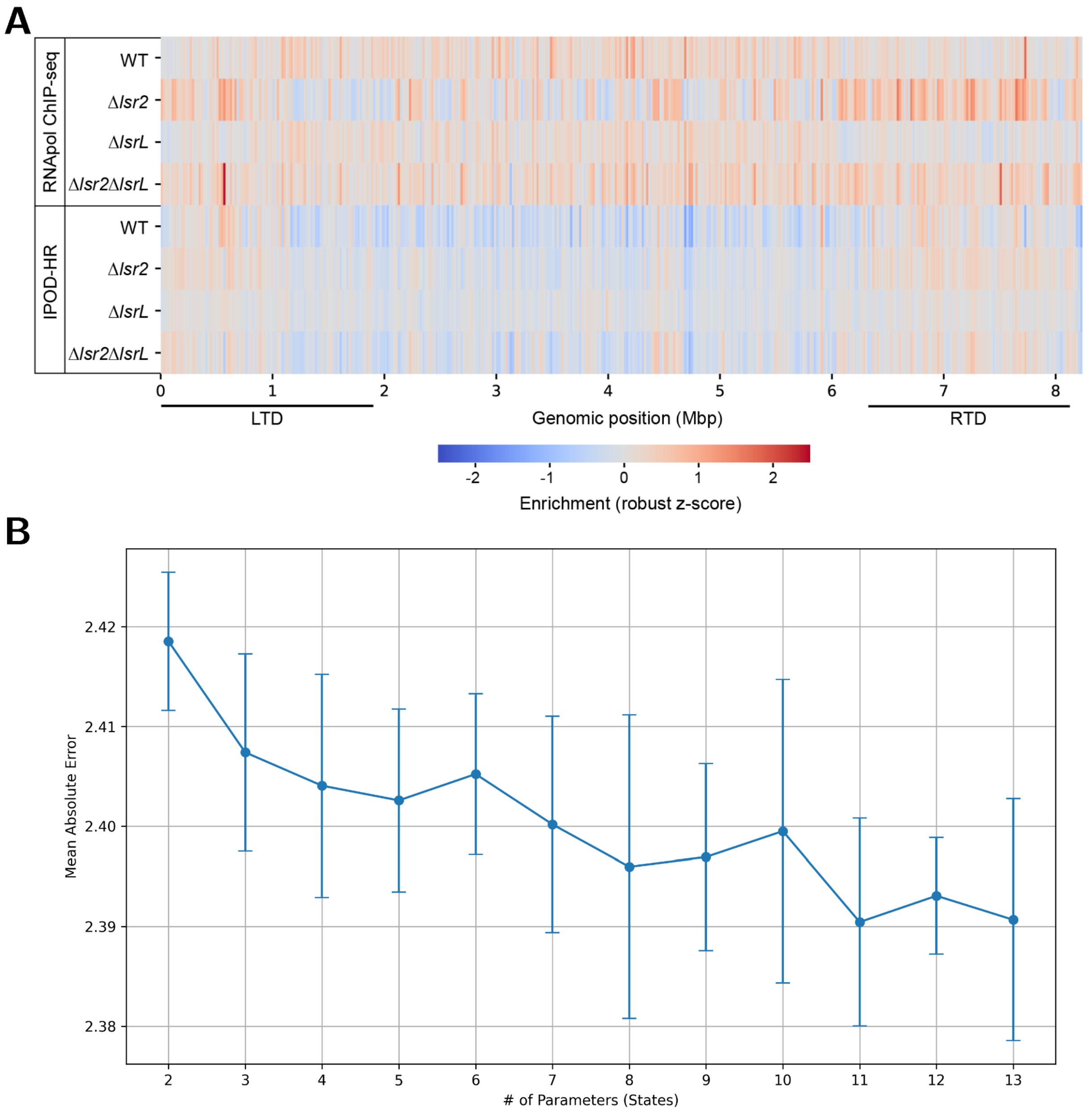


**Supplementary Figure 1. Genome-wide protein occupancy and 5-fold cross-validated mean absolute error of the GHMM state selection.** (**A**) Heatmap of the DNA-binding protein occupancy derived from RNApol ChIP-seq (rows 1-4) and IPOD-HR (rows 5-8) in the WT, Δ*lsr2*, Δ*lsrL*, and Δ*lsr2*Δ*lsrL* mutants across the chromosome. Enrichment is represented as robust z-scores (relative to input) that were calculated as rolling averages in 20 kb windows. The LTD and RTD, separated by the core region (~1.9-6.3 Mbp), are indicated. (**B**) The 5-fold cross validation (CV) was conducted on a GHMM using the RNApol ChIP-seq, IPOD-HR, and Log2 TMM-normalized RNA-seq coverage from the WT, Δ*lsr2*, Δ*lsrL*, and Δ*lsr2*Δ*lsrL* mutants, with states ranging from 2 to 13. The 5-fold cross-validation was performed to select the optimal number of states for inference of chromatin states. The error bars indicate 95% confidence interval for the mean absolute error for each state, computed as 1.96 times the standard error of the mean.

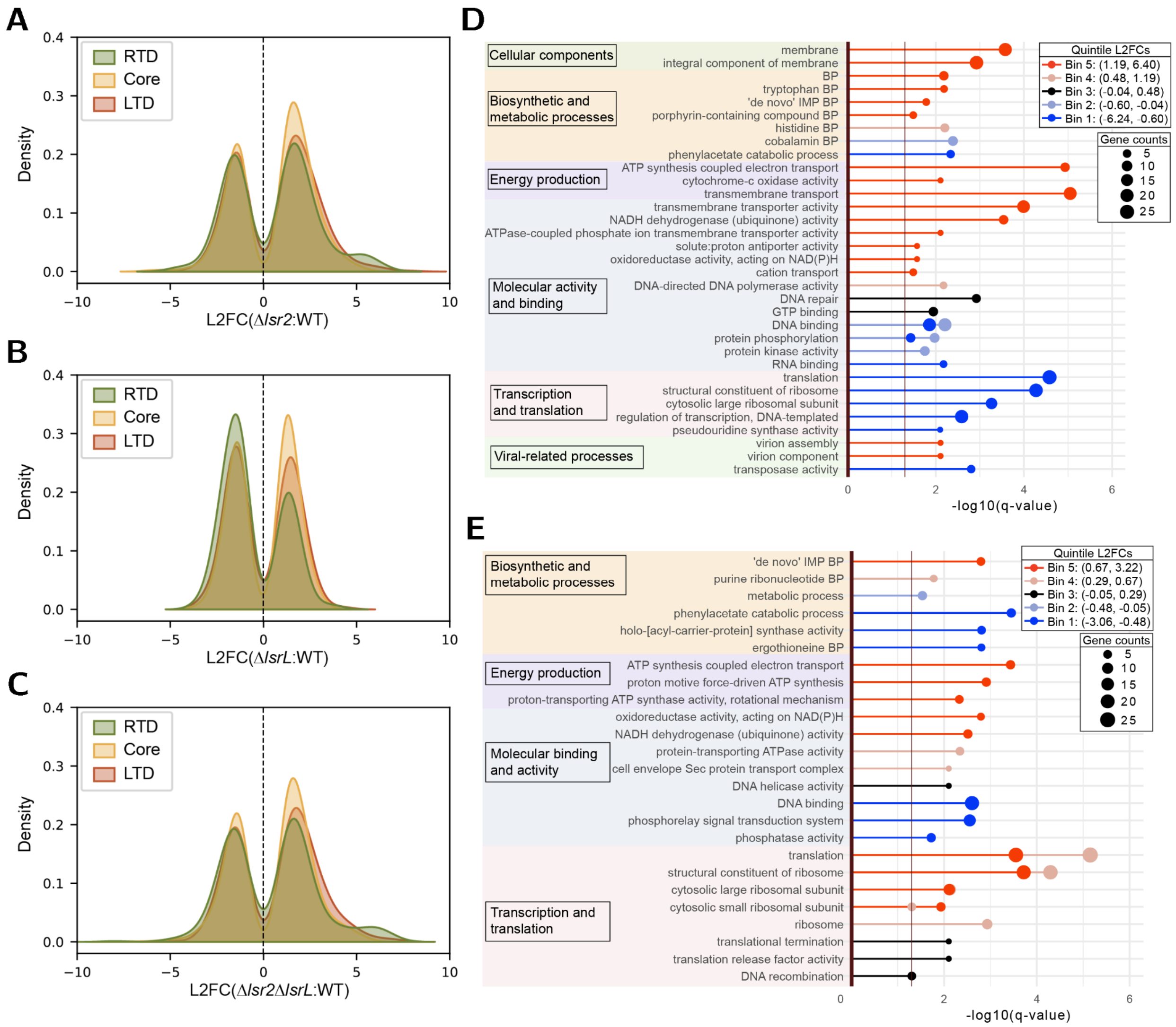


**Supplementary Figure 2. Genome-wide expression profiles and GO term enrichments highlight regional and regulatory differences across mutants.** (**A**-**C**) Kernel density distributions of significant gene expression changes (|L2FC| ≥ 1, q-value < 0.05) for the (**A**) Δ*lsr2*, (**B**) Δ*lsrL*, and (**C**) Δ*lsr2*Δ*lsrL* mutants (relative to WT). Curves are grouped by chromosomal region: LTD (0–1.9 Mbp; red), core (1.9–6.3 Mbp; orange), and RTD (6.3–8.22 Mbp; green). Densities are weighted so that the area under each region's curve reflects its share of all significant genes. (**D**-**E**) Lollipop plots displaying the GO term enrichment analysis of differentially expressed genes in (**D**) Δ*lsr2* and (**E**) Δ*lsrL* mutants. The color of the lollipop heads and sticks indicate the GO terms that are highly upregulated or downregulated relative to WT according to the L2FCs representing each bin (see key for details). Each lollipop head is centered on its corresponding x-axis position, with the length of the head and sticks representing the significance (-log10(q-value)). GO terms including 'BP' refer to biosynthetic processes and IMP refers to inosine monophosphate.

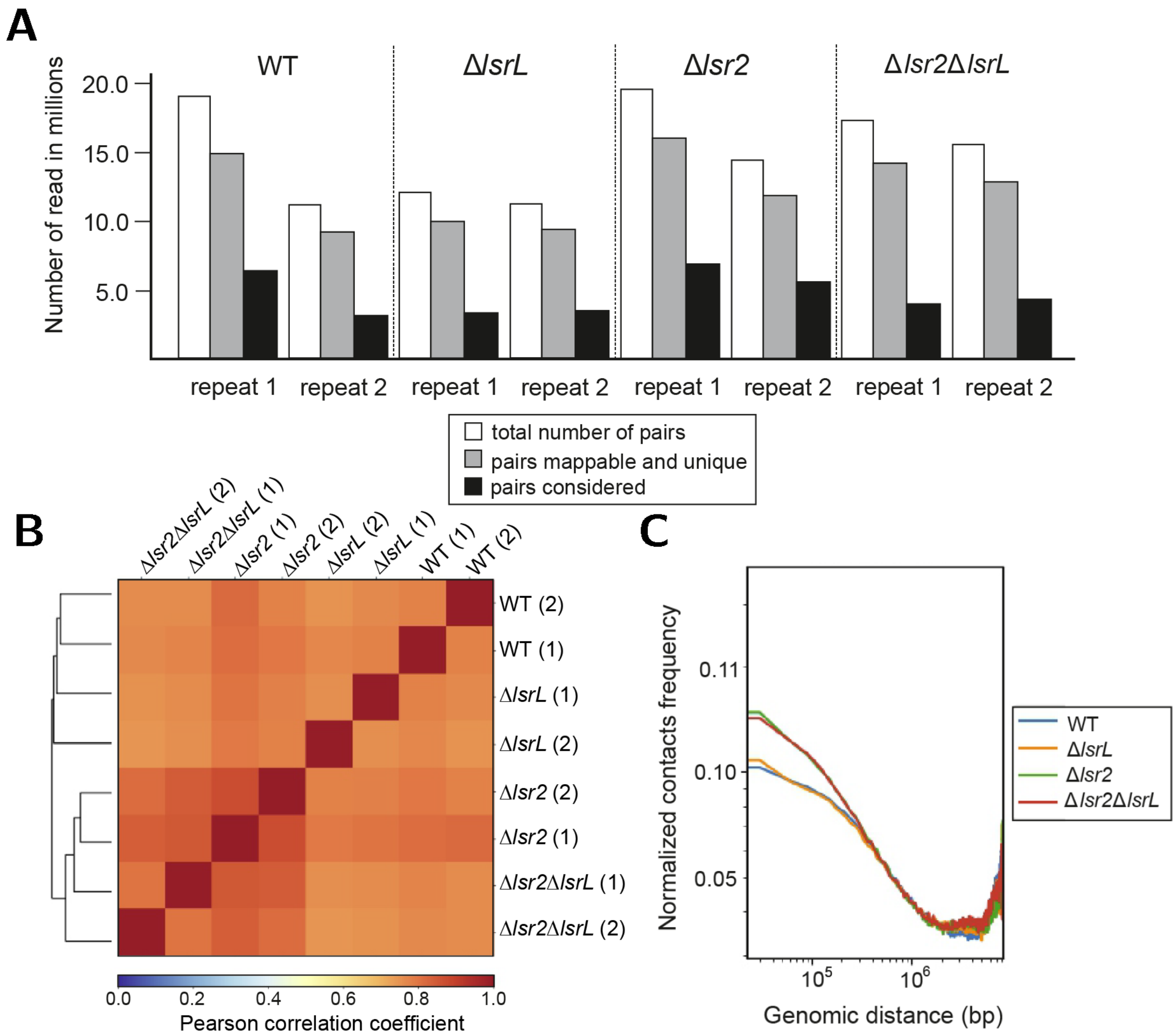


**Supplementary Figure 3. Statistics and data reproducibility of Hi-C**. (**A**) Summary of raw sequencing data for Hi-C matrix preparation for particular *S. venezuelae* strains (WT, Δ*lsrL*, Δ*lsr2*, and Δ*lsr2*Δ*lsrL*). The diagram shows the number of paired reads Δ*lsr2* (1) (white), the number of pairs mapped to the chromosome (gray) as well as the number of pairs successfully filtered and used for Hi-C matrix construction (black). Each analysis was performed in two biological replicates (replicate #1 and replicate #2). (**B**) Pairwise correlation analysis of the normalized Hi-C matrices for indicated *S. venezuelae* strains. The Hi-C matrices were grouped by their similarities based on the Pearson correlation coefficient. The biological replicates are numbered (1 or 2). (**C**) The normalized Hi-C contacts frequency between two DNA loci plotted in relation to genomic distance.

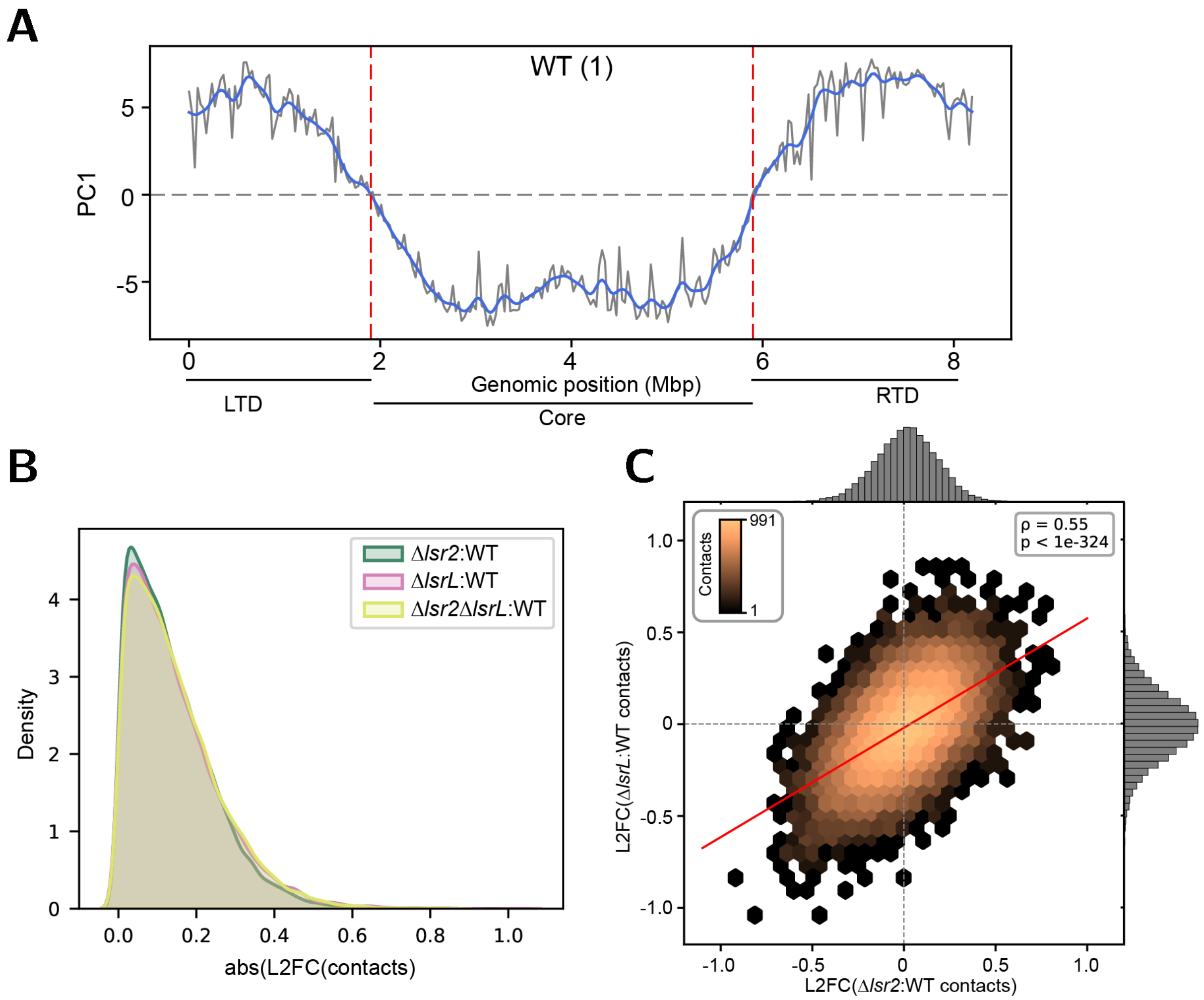


**Supplementary Figure 4. Identification of chromosomal regions and comparison of Hi-C contact profiles across mutants.** (**A**) Principal component 1 (PC1; gray) from principal components analysis of the WT Hi-C contact matrix (replicate 1) plotted across the chromosome. PC1 values were smoothed using a Gaussian filter with $\sigma$ = 2 bins, corresponding to a smoothing window of ~60 kb given the 30 kb resolution of the Hi-C map (blue). Red dashed lines indicate the boundaries of the LTD (~0-1.9 Mbp) and RTD (~5.9-8.22 Mbp). (**B**) Distributions of absolute L2FC of Hi-C contacts for the Δ*lsr2* (green), Δ*lsrL* (pink), and Δ*lsr2*Δ*lsrL* (yellow) mutants relative to WT. (**C**) Hexbin plot comparing L2FC of Hi-C contacts between the Δ*lsr2* and Δ*lsrL* mutants (both relative to WT). The plot shows the density of contacts (log-scaled) and a linear regression line (slope = 0.59). The Pearson correlation coefficient and corresponding p-value between the two differential contact profiles is displayed.

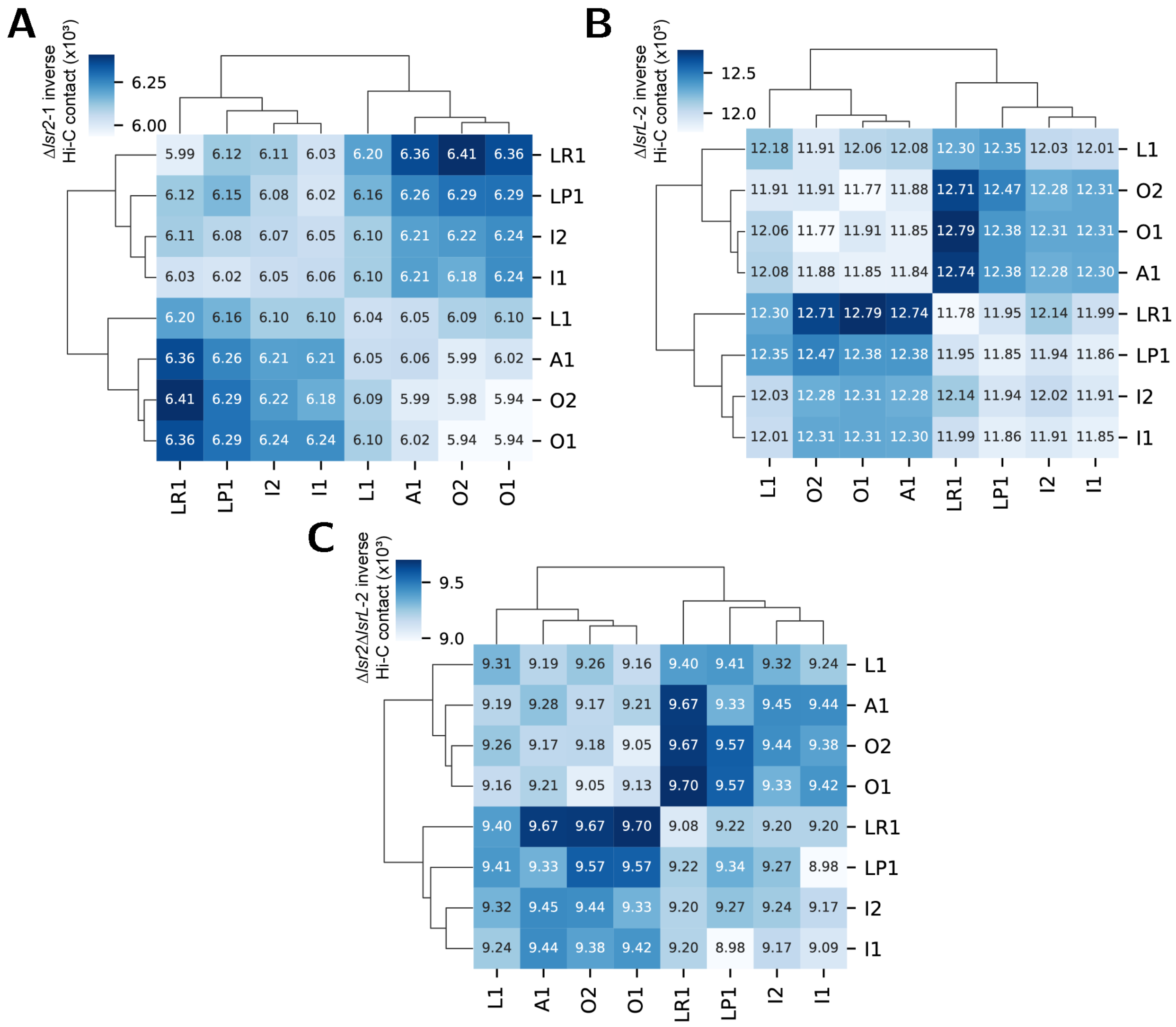


**Supplementary Figure 5. Hierarchical clustering of median Hi-C distances between chromatin states in Δ*lsr2*, Δ*lsrL*, and Δ*lsr2*Δ*lsrL* mutants, reflecting 3D spatial organization.** (**A-C**) Heatmaps of median spatial distances between chromatin state pairs inferred from (**A**) Δ*lsr2* replicate 1, (**B**) Δ*lsrL* replicate 2, and (**C**) Δ*lsr2*Δ*lsrL* replicate 2 Hi-C contact matrices at a 30 kb bin resolution. Replicates were selected based on having the fewest bins with zero Hi-C contacts. Distances are computed as the log10-transformed inverse Hi-C contact (x$10^3$) between bins assigned to each chromatin state, excluding local interactions (± 750 kb). Lower inverse Hi-C contacts values indicate closer spatial proximity, while higher values reflect greater spatial proximity between chromatin states. States are clustered by Manhattan distance to reveal 3D spatial compartmentalization.

### Supplemental Tables

**Supplementary Table 1. AntiSMASH predicted BGC clusters within *S. venezuelae*.** BGC types were clustered into a broader BGC class based on the general type of specialized metabolites they are predicted to produce. Provided in the attached “Supplementary Tables” file.

**Supplementary Table 2. GHMM transition state probabilities between inferred chromatin states.** The transition state probabilities between chromatin states in the 9-state model trained on the RNApol ChIP-seq, IPOD-HR, and normalized RNA-seq read coverage from the WT, Δ*lsr2*, Δ*lsrL* and Δ*lsr2*/Δ*lsrL* mutants. Provided in the attached “Supplementary Tables” file.
